# Oscillatory movement of a dynein-microtubule complex crosslinked with DNA origami

**DOI:** 10.1101/2021.12.06.471346

**Authors:** Shimaa A. Abdellatef, Hisashi Tadakuma, Kangmin Yan, Takashi Fujiwara, Kodai Fukumoto, Yuichi Kondo, Hiroko Takazaki, Rofia Boudria, Takuo Yasunaga, Hideo Higuchi, Keiko Hirose

## Abstract

During repetitive bending of cilia and flagella, axonemal dynein molecules move in an oscillatory manner along a microtubule (MT), but how the minus-end-directed motor dynein can oscillate back and forth is unknown. There are various factors that may regulate the dynein activities, e.g., the nexin-dynein regulatory complex, radial spokes, and central apparatus. In order to understand the basic mechanism of the oscillatory movement, we constructed a simple model system composed of MTs, outer-arm dyneins, and DNA origami that crosslinks the MTs. Electron microscopy (EM) showed patches of dynein molecules crossbridging two MTs in two opposite orientations; the oppositely oriented dyneins are expected to produce opposing forces. The optical trapping experiments showed that the dynein-MT-DNA-origami complex actually oscillate back and forth after photolysis of caged ATP. Intriguingly, the complex, when held at one end, showed repetitive bending motions. The results show that a simple system composed of ensembles of oppositely oriented dyneins, MTs, and inter-MT crosslinkers, without the additional regulatory structures, has an intrinsic ability to cause oscillation and repetitive bending motions.

## Introduction

Beating of cilia and flagella is powered by axonemal dynein molecules that are minus-end-directed MT motors. Each dynein is tethered to a doublet MT with its tail and interacts with the neighboring MT using its stalks in an ATP dependent way, to produce relative sliding of the two MTs. The MTs in an axoneme are arranged in a 9+2 structure, interconnected with various components, such as the nexin-dynein regulatory complex (Heuser *et al*, 2009). Because the neighboring doublet MTs are interlinked and also anchored to the basal body at the base, minus-end directed movement of the dynein molecules on one side of the axoneme causes bending of the axoneme in one direction. During the bending, the dyneins on the opposite side of the axoneme are pulled toward the MT plus end. Thus, during cyclical bending of the axoneme, dynein molecules should oscillate back and forth along a MT. How a minus-end-directed motor dynein can oscillate along a MT is unknown.

There are many factors that may influence the force production of dyneins. For example, outer-arm dyneins and several species of inner-arm dyneins are thought to affect each other’s activities. The arrangement of the dyneins in an axoneme must be also important for their communication; the outer-arm dyneins are aligned in a row with a periodicity of 24 nm, and the inner-arm dyneins are also regularly placed in specific positions. The inter-doublet linkers connect the circumferentially arranged nine doublet MTs, and thus transmit the force locally produced by some dyneins to the other parts of the axoneme. Other regulatory components, such as the radial spokes and central apparatus also seem to play important roles in the regulation (Witman *et al*, 1978). In order to avoid the complexity and understand the basic mechanism of cilia and flagella motility, we designed a simple model system and tested its structure and motility. Our system consists of *in vitro* polymerized MTs, axonemal outer-arm dynein molecules and passive linkers that interconnect a pair of MTs (Fig 1A). This simple model system does not have the circular arrangement of nine MTs in the axoneme, nor does it have inner-arm dyneins, radial spokes or the central apparatus.

**Figure 1.**
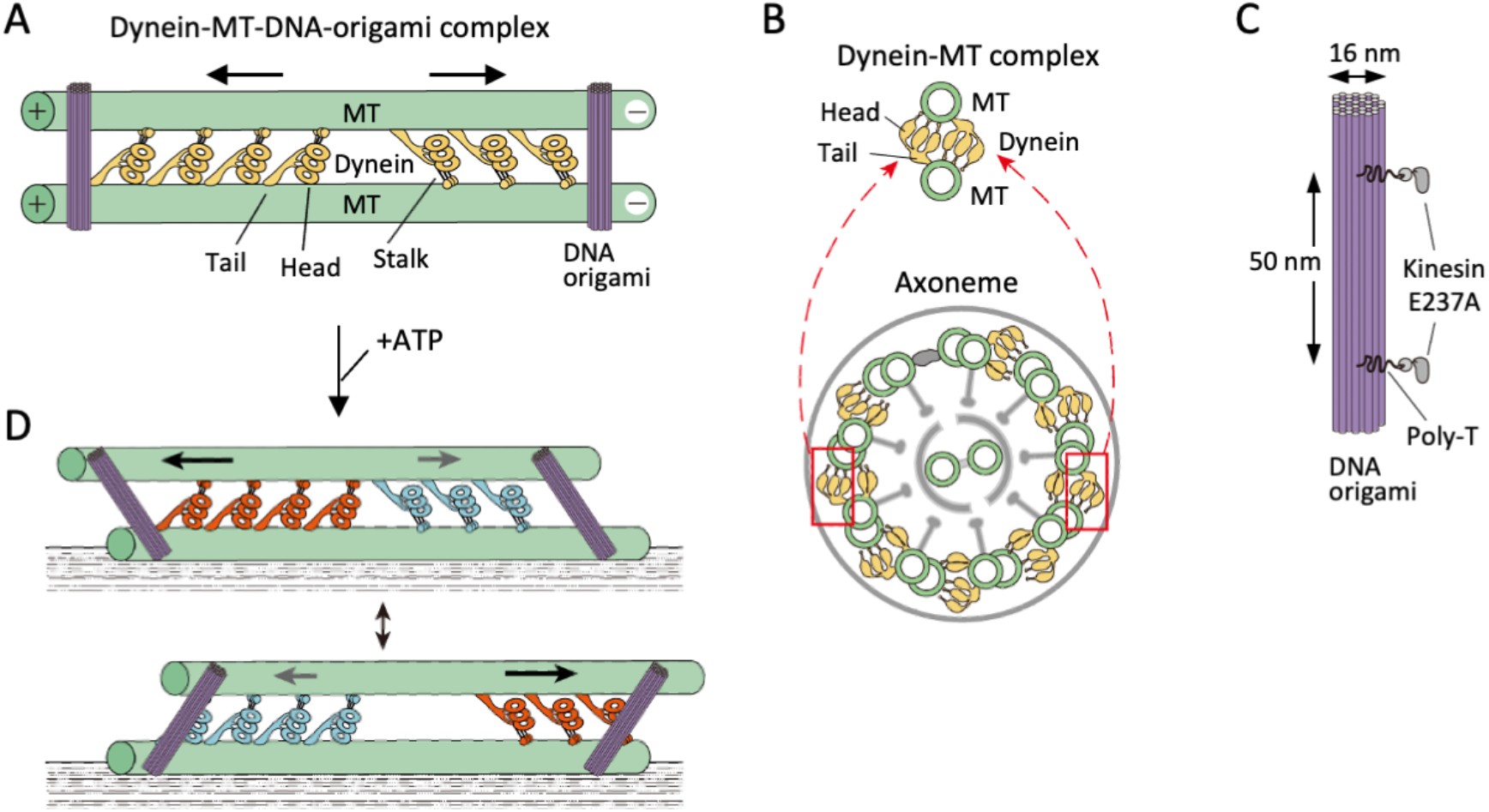
Design and model for motility of the dynein-MT-DNA-origami complex. (A) Geometry of the dynein-MT-DNA complex. *Chlamydomonas* outer-arm dynein molecules crossbridge 2 MTs, binding with their tails and stalks. The stalks point toward the MT minus end. Because the tails and stalks bind to different MTs, there are two possible orientations for dynein depending on which MT the stalks bind to. The two orientations of dynein molecules produce force in opposite directions (indicated by arrows). (B) Schematic illustration of a cross-sectional view of the dynein-MT complex in comparison with that of an axoneme. Note that the geometry of dyneins in the dynein-MT complex mimics that of a combination of the two boxed regions of an axoneme. (C) Design of rod-shaped DNA origami. For binding to MTs, mutant kinesin motor domains (E237A) were attached to 5’-SNAP-ligand-modified handles placed on the DNA rod. The two handles were separated by ~50 nm, corresponding to the center-to-center distance of the two MTs in the dynein-MT complex. Each handle has 30 nucleotides of poly-thymidine linker between the SNAP ligand and the rod. (D) A model showing oscillatory movement of a dynein-MT-DNA-origami complex. DNA rods are expected to crosslink the two MTs of the complex and restrict their relative movement. If the two groups of dynein molecules alternately produce higher forces, the complex is expected to show oscillatory back and forth movement.

To reconstitute the regular arrangement of outer-arm dyneins, which allows neighboring dynein molecules to interact with each other, we used dynein preparations extracted from *Chlamydomonas* flagella axonemes with high salt (typically 0.6 M KCl). The high-salt-extracted dynein preparations are known to crossbridge MTs with ~24-nm periodicity (Haimo *et al*, 1979), and although they contain other components in addition to outer-arm dyneins, previous work showed that the proteins bound to MTs are mostly outer-arm dynein (Aoyama & Kamiya, 2010; Oda *et al*, 2007). As inter-MT linkers, we utilized rod-shaped DNA origami structures and crosslinked the MTs *via* flexible linkers (Fig 1C). Use of DNA origami enabled us to design a molecular layout of the crosslinking structure that allows relative sliding of the MTs over a certain distance, resulting in oscillation and repetitive bending motions as seen in intact axonemes.

## Results

### Geometry of Dynein-MT complexes

Dynein preparations extracted with high salt from *Chlamydomonas* flagella axonemes were mixed with taxol-stabilized MTs polymerized from brain tubulin. As previously reported (Aoyama & Kamiya, 2010; Haimo *et al*., 1979; Oda *et al*., 2007), the MTs became bundled with the crossbridging molecules bound with a period of ~24 nm (Figs 2A and 2B). When a low concentration of the dynein preparation (6.25–12.5 µg/ml (corresponding to ~3-6 nM outer-arm dynein)) was mixed with 20-25 µg/ml MTs (200-250 nM tubulin dimers), most of the MT bundles were thin, composed of 2 to several MTs, so that in many cases we were able to observe a single layer of crossbridges without overlapping. Negative-stain EM observation of the individual crossbridges showed characteristic shapes of outer-arm dynein, with stacked heads and a tail (Heuser *et al*., 2009; Movassagh *et al*, 2010) (Fig 2C), confirming that the crossbridging proteins are mostly outer-arm dyneins. The average number of the dyneins per 1 µm of a MT pair was about 12 when 6.25 µg/ml dynein was used.

**Figure 2.**
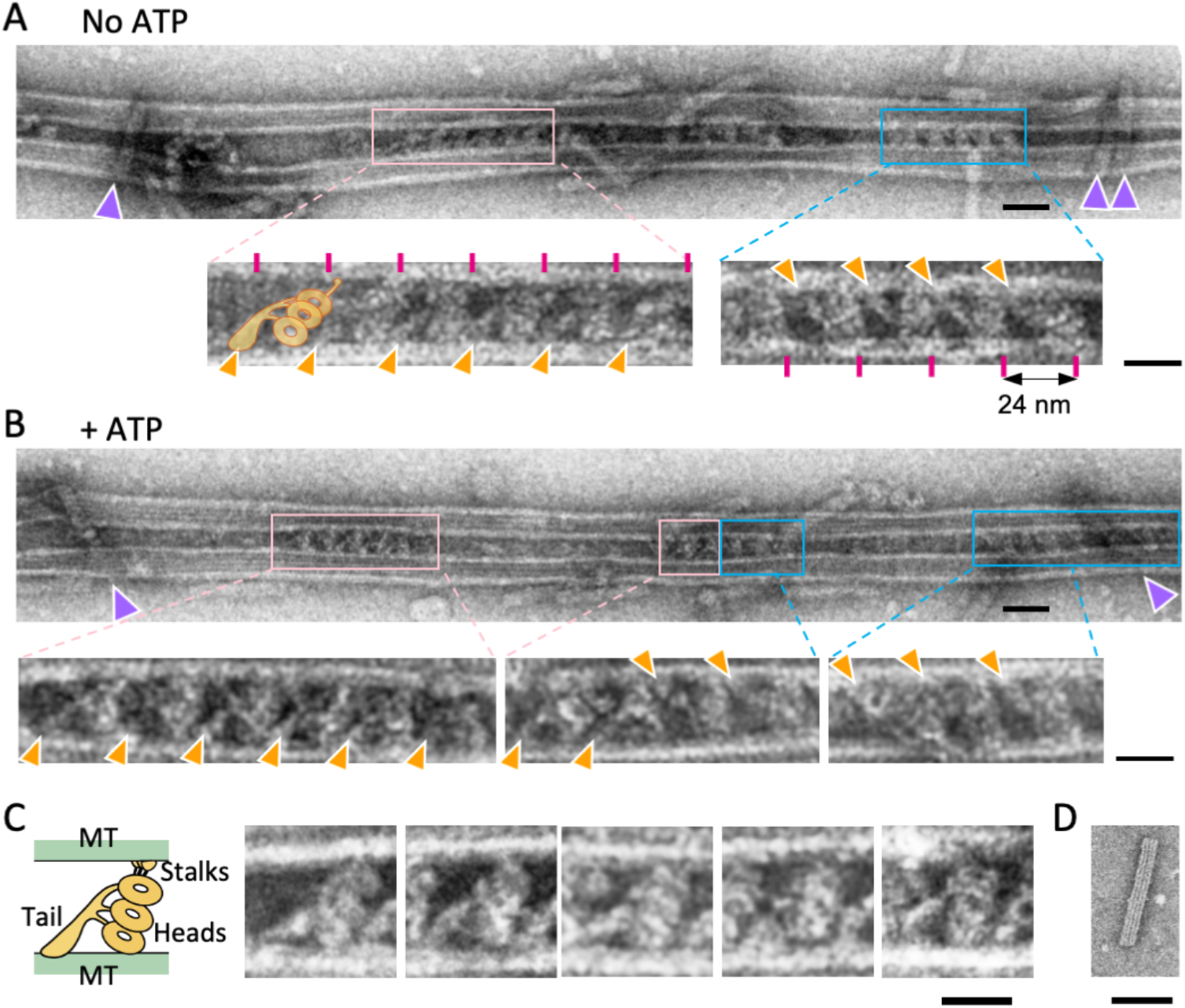
Structure of the dynein-MT-DNA-origami complex. (A, B) Negative stain images of the dynein-MT-DNA-origami complex in the absence (A) and presence (B) of 0.1 mM ATP. DNA rods crosslinking the MTs are indicated by purple arrowheads. Dynein molecules tend to bind in patches and the neighboring dyneins usually have the same orientation. Patches of dynein cross-bridging the MTs in two different orientations are indicated by pink and cyan boxes. Enlarged views show examples of dyneins in two orientations, with their tails (indicated by orange arrowheads) on different MTs. Magenta lines show a periodicity of 24 nm. The typical shape of an outer-arm dynein is illustrated. Bars, 50 nm in low magnification images and 20 nm in enlarged views. (C) Images of individual dynein molecules crossbridging 2 MTs. As illustrated on the left, the characteristic shapes of a *Chlamydomonas* outer-arm dynein molecule (Heuser *et al*., 2009; Movassagh *et al*., 2010) with a tail and three heads are observed. Bar: 20 nm. (D) A negative stain image of a DNA origami rod. Bar: 50 nm.

When a dynein molecule crossbridges a pair of MTs, its tail is fixed onto one of the MTs and the stalks interact with the other MT in a nucleotide-dependent manner. Therefore, depending on which MT the tail binds to, there are two possible orientations (Fig 1A). EM images confirmed that dyneins crossbridge MTs in two different orientations (Fig 2). As observed previously with sea urchin dynein (Hirose, 2012; Ueno *et al*, 2008), the crossbridging dyneins made patches, and the adjacent dynein molecules within one patch tended to have the same orientation. *In vivo*, outer-arm dynein binds to MTs with its head+stalk oriented toward the minus end of the MTs, and its tail oriented toward the plus end (Goodenough & Heuser, 1982; Heuser *et al*., 2009; Movassagh *et al*., 2010), so that the two MTs crosslinked with dyneins have the same polarity (Fig S1A). We have previously shown by cryo-EM analysis that *in vitro* also, dynein’s head and stalk are always oriented toward the minus end of the MT to which its stalk binds (Ueno *et al*., 2008). Thus, we determined the polarity of the MTs in the dynein-MT complex based on the orientation of dyneins in the EM images. Although the two MTs of a complex were often antiparallel in the case of sea urchin dynein (Ueno *et al*., 2008; Yokota & Mabuchi, 1994), the majority of the *Chlamydomonas*-dynein-MT complexes (36 out of 44 complexes) showed parallel arrangements of the MTs as in axonemes (Figs S1B and S1C). In the complex with parallel MTs, the dynein molecules in two opposite orientations are expected to produce force in opposite directions (Figs 1A and S1A). This arrangement, which is similar to the models previously proposed (Camalet & Jülicher, 2000; Mitchison & Mitchison, 2010; Riedel-Kruse *et al*, 2007), is analogous to a combination of two dynein arrays on the opposite sides of an axoneme (Fig 1B), which produce antagonistic forces. The resemblance in their geometries led us to the idea that the dynein-MT complex might work as a simple model system to investigate the minimum components required for oscillatory movements that occur in cilia and flagella.

### Motile properties of the dynein-MT complex

The outer-arm dynein preparations used here supported movement of MTs in gliding assays at a speed comparable to those in previous work (Alper *et al*, 2013; Furuta *et al*, 2009) (Fig S2A). We then tested the motility of dynein-MT complexes that contain a pair of MTs crossbridged by dyneins in two different orientations (Fig S2B). When the dynein-MT complexes are adsorbed to a glass surface, the MTs that are directly attached to the glass cannot move, but other MTs in the same complex are allowed to slide relative to the glass-attached MTs. Since addition of ATP causes relative sliding of the MTs and disassembly of the complex, we used caged ATP and observed the movement of MTs immediately after photolysis of caged ATP. Even though the majority of the complexes are thought to contain groups of dynein molecules that produce force in opposite directions, many of the complexes showed unidirectional sliding of MTs (Movie S1), probably because the numbers of molecules in the two groups are not equal and the major group would ‘win’. The velocity was variable (Fig S2B), with some MTs sliding much faster than those observed in gliding assays. Fast sliding was previously observed with similar dynein-MT complexes but with much higher concentration of dynein, and was interpreted as the effect of cooperation of dyneins aligned with the 24-nm periodicity (Aoyama & Kamiya, 2010).

We have also measured the force produced by the dynein-MT complexes using optical trapping methods (Fig S3). The average number of dyneins that crossbridge the pair of MTs was estimated to be ~35 from inspection of the EM images (12 molecules/µm) and the average length of the MT (2.9 ± 0.9 μm (mean ± sd)). Typical traces showed a maximum force of ~30 pN. The value is several times larger than the force produced by single molecules of outer-arm dynein from sea urchin (5-6 pN) (Shingyoji *et al*, 2015) or *Tetrahymena* (4.7 pN) (Hirakawa *et al*, 2000), indicating cooperative force production. On the other hand, it is smaller than the simple sum of the forces produced by each dynein. Dependence of the maximum force on the number of motors differs in different motors (Furuta *et al*, 2013; Soppina *et al*, 2009). The partial dependence observed here for outer-arm dynein may be related to the weak processivity of the molecule. Alternatively, the dynein molecules oriented in the opposite direction may act as a load.

### Crosslinking of the dynein-MT complex with DNA origami structures

The above results showed that the dynein-MT complex alone is not sufficient to generate oscillatory movement, even though it contains two groups of dyneins that produce force in opposite directions: the movement was mostly in one direction and the complex disassembled. To prevent disassembly and mimic axonemal doublet MTs that are interconnected with linkers, we crosslinked the MTs of the dynein-MT complex. A rod-shaped DNA origami of ~84 nm in length (Fig 1C and Table S1) was attached to the dynein-MT complex via immotile mutant kinesin motor domains (E237A) (Rice *et al*, 1999). Flexible poly-T linkers were inserted between the DNA rod and kinesin to allow tilting of the rod and relative sliding of the MTs up to ~200 nm (Fig S4A). *In vitro* motility assays confirmed that addition of enough DNA origami rods (e.g. 2.5 nM DNA rods for the dynein-MT complex with 6.25 µg/ml dynein) can actually prevent disassembly of the complex in the presence of ATP (Movie S2).

DNA origami rods crosslinking the dynein-MT complex were clearly visible by EM (Fig 2). The average number of DNA rods crosslinking a pair of MTs was ~2.4 per µm when 2.5 nM DNA rods were used. As expected, the binding angle in the absence of ATP was variable but centered around the perpendicular (Fig S4C). When ATP was added to the complex, the deviation of the rod angle from the perpendicular increased (p<0.002, Mann-Whitney U-test), indicating that the DNA rods tilted because of relative sliding of the MTs.

### Dynein-MT-DNA-origami complexes show oscillatory movements

We then investigated the motility of the dynein-MT-DNA-origami complexes in detail. Since the expected relative sliding of the MTs was too small to be detected by fluorescence microscopy, we attached a bead to the complex and measured the displacement after photolysis of caged ATP (Fig 3). In some cases, the bead bound to a MT moved unidirectionally and then abruptly went back to the trap center, probably because of detachment of dynein from the MT (e.g., the first ~100 ms after the UV flash in Fig 3B), as observed in previous work using single dynein molecules (e.g., (Hirakawa *et al*., 2000; Sakakibara *et al*, 1999; Toba *et al*, 2006)).

**Figure 3.**
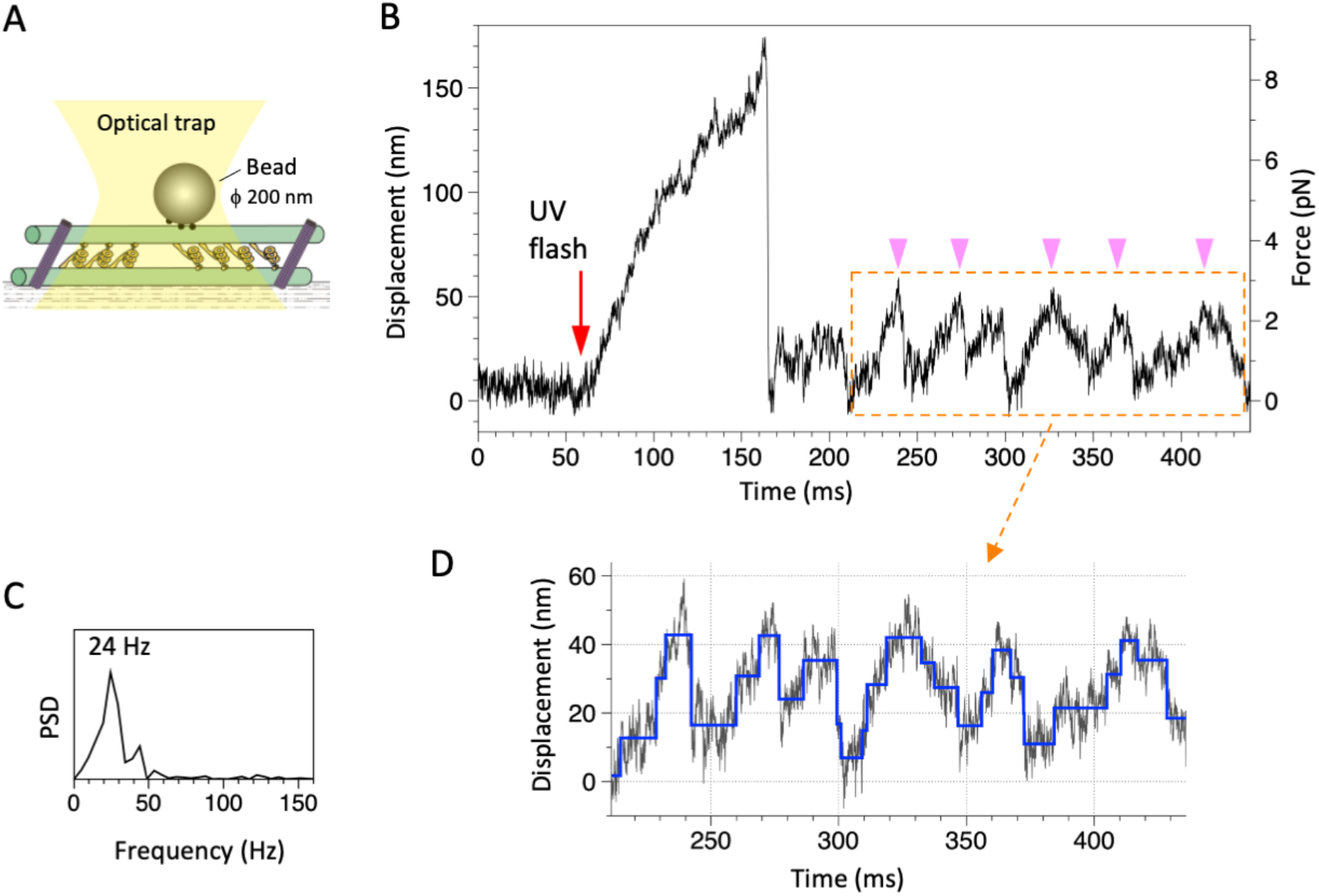
Movement of the dynein-MT-DNA-origami complex measured in optical trapping assays. (A) Schematic of the experimental set-up (not to scale). A streptavidin-coated bead was captured by the optical trap (trap stiffness 0.052 pN/nm) and attached to a dynein-MT-DNA-origami complex adsorbed to a glass surface. (B) Trace showing displacement of a bead attached to the dynein-MT-DNA-origami complex after UV photolysis of caged ATP (red arrow). A part of the trace shows oscillatory movement (pink arrowheads). (C) The frequency of the oscillatory movement was measured by the power spectral density (PSD). (D) Steps (blue) detected for the same region by the step-fitting algorithm (Kerssemakers *et al*., 2006). Steps are found in both forwards (away from the trap center) and backwards (toward the trap center) movements.

The most notable feature observed in the optical trapping measurement of the dynein-MT-DNA-origami complex was oscillatory movements. Typical traces are shown in Figs 3B, 3D, and S5. Although the oscillation was irregular, it was clearly different from the noisy vibration before the UV flash (compare the trace before the UV flash and the region boxed in orange in Fig 3B), and the power spectrum density showed clear peaks (Figs 3C and S5, inset), which were not observed before the UV flash. Since caged ATP was locally photolyzed using a UV spot with a full width at half maximum of ~20 µm and the released ATP diffuses in the solution, we analyzed the traces typically within ~0.5 s after photolysis of ATP. Forty-eight out of 94 such traces showed sliding movement, and 75% of them exhibited oscillatory movement in some parts of the trace. The amplitude of the oscillatory movements was variable, averaging 26.6 nm and 26.4 nm for the forward (away from the trap center) and backward (back toward the trap center) movement, respectively (Fig 4B). The amplitudes much smaller than the expected maximum sliding distance (~200 nm) can be explained by the fact that this maximum distance is achieved only when all the DNA rods bind to the MTs at the ideal tubulin subunits and in a perpendicular angle (Figs S4A and S4B): since the MTs are crosslinked with multiple DNA linkers with various angles (Fig S4C), the actual sliding distance is thought to be significantly shorter than the maximum sliding distance. The velocities of the forward and backward movements during oscillation were also variable (Fig 4C), but they were both in the same range as the velocity observed in the gliding assay of the dynein-MT complex without DNA rods (4.4 µm/s in average; Fig S2B). The difference between the averaged velocities of the forward and backward movements (3.8 µm/s and 6.4 µm/s, respectively)) may be because the force of the optical trap works as a load for the forward movement, whereas it assists the backward movement. Some detachment may also contribute to the faster velocity of the backward movement. The frequencies of the oscillation were measured using the power spectrum of the traces (Figs 3C and S5). The frequencies were variable, with an average of 32.9 Hz (Fig 4D).

**Figure 4.**
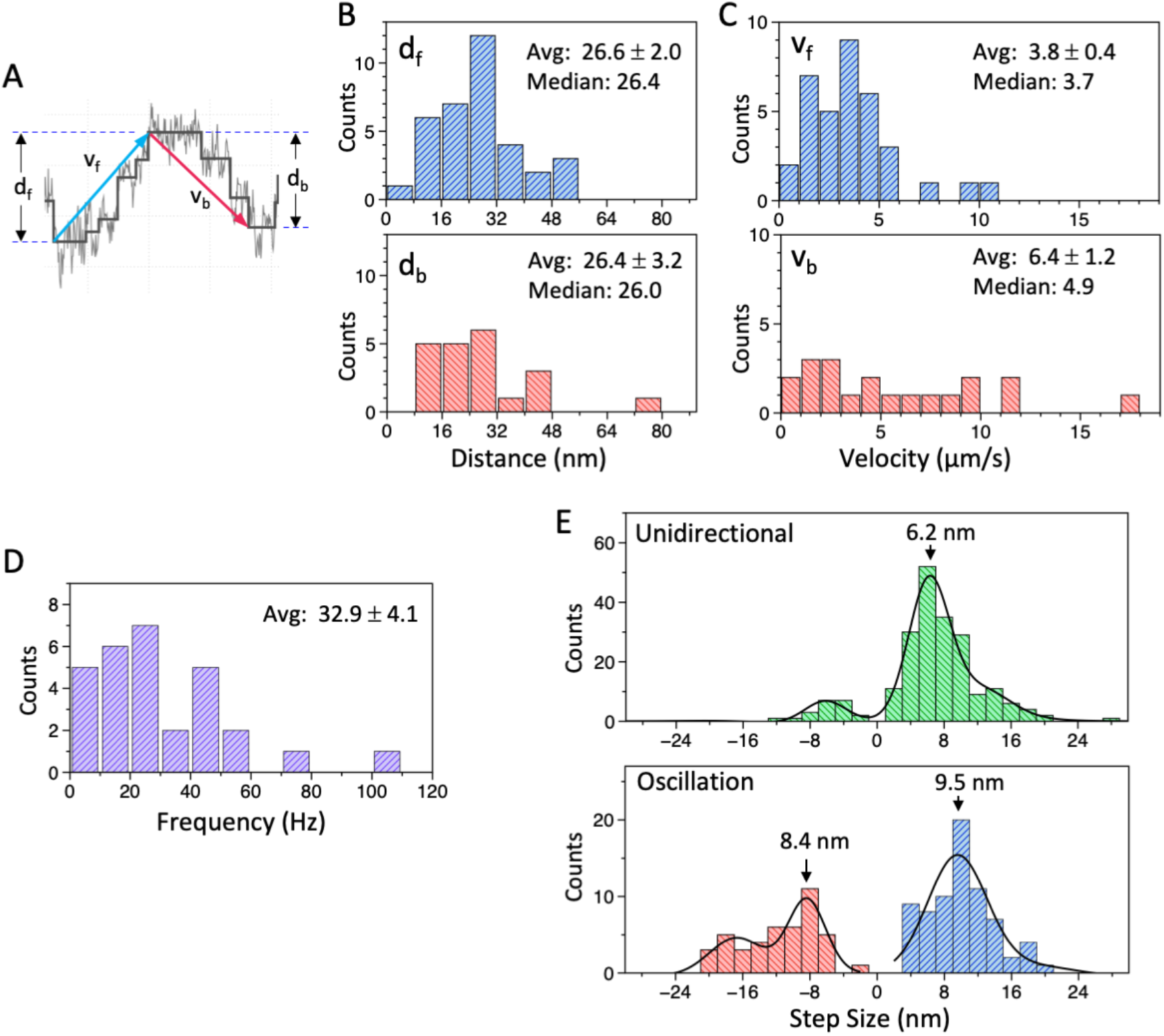
Analysis of movement of the dynein-MT-DNA-origami complex measured in optical trapping assays. (A) Definition of the distance (d_f_, d_b_) and velocity (v_f_, v_b_) used for the histograms in (B) and (C). (B, C) Histograms of the distance (B) and velocity (C) during the forwards (d_f_, v_f_) and backwards (d_b_, v_b_) movements of the bead attached to the dynein-MT-DNA-origami complex. Average (mean ± s.e.m.) and median values are indicated in each histogram (N = 35, 21, 35, 21 for d_f_, d_b_, v_f_, v_b_, respectively). (C) Histograms of the frequency of the oscillatory movements (N = 29). (D) Histograms of the step size during unidirectional movement (top; N=211) and oscillatory movement (bottom; N=72 for forwards and N=44 for backwards), fit with multiple Gaussian curves. Main peak positions of the Gaussian functions are indicated.

Although many of the traces were noisy, a step-finding algorithm (Kerssemakers *et al*, 2006) detected stepwise movements in both the unidirectional and oscillatory movements of some traces (Fig 3D). Previous work using single *Tetrahymena* dynein molecules reported steps of ~8 nm at an extreme condition: very low concentration of ATP (3 µM) (Hirakawa *et al*., 2000). Cytoplasmic dynein also showed predominantly 8 nm steps (Reck-Peterson *et al*, 2006; Toba *et al*., 2006), but the step size depended on load (Belyy *et al*, 2014; Gennerich *et al*, 2007). Although many of the traces of our dynein-MT-DNA-origami complex were noisy and we could not accurately measure the step sizes, the peaks of the step sizes were close to 8 nm (6.2 nm, 8.4 and 9.5 nm for unidirectional movement, forward, and backward displacements of the oscillatory movement, respectively) (Fig 4E). If the oscillatory movements we observed are simply the cycles of unidirectional movement and dissociation of dyneins, we would not detect multiple steps during the backward movements. Thus, the results indicate that both forward and backward displacements during the oscillatory movement are steps of dynein.

### An ensemble of uniformly-oriented dynein molecules moves a MT unidirectionally

The above results show that a dynein-MT complex that contains two groups of oppositely oriented dyneins can move in an oscillatory manner when the MTs are connected with DNA origami linkers, whereas a dynein-MT complex without the linkers does not oscillate. However, previous work using single or a few molecules of outer-arm dynein also reported bi-directional movements over a range of several tens of nanometers (Shingyoji *et al*, 1998; Shingyoji *et al*., 2015). We thus examined whether the oscillatory movement was also observed with a MT interacting with multiple dyneins but in the absence of oppositely oriented dyneins. Two experimental designs were used: optical trapping measurements of MTs gliding over dynein-coated surfaces (Fig 5A), and dynein-MT complexes in which all the dyneins are in the same orientation (Fig 5B).

**Figure 5.**
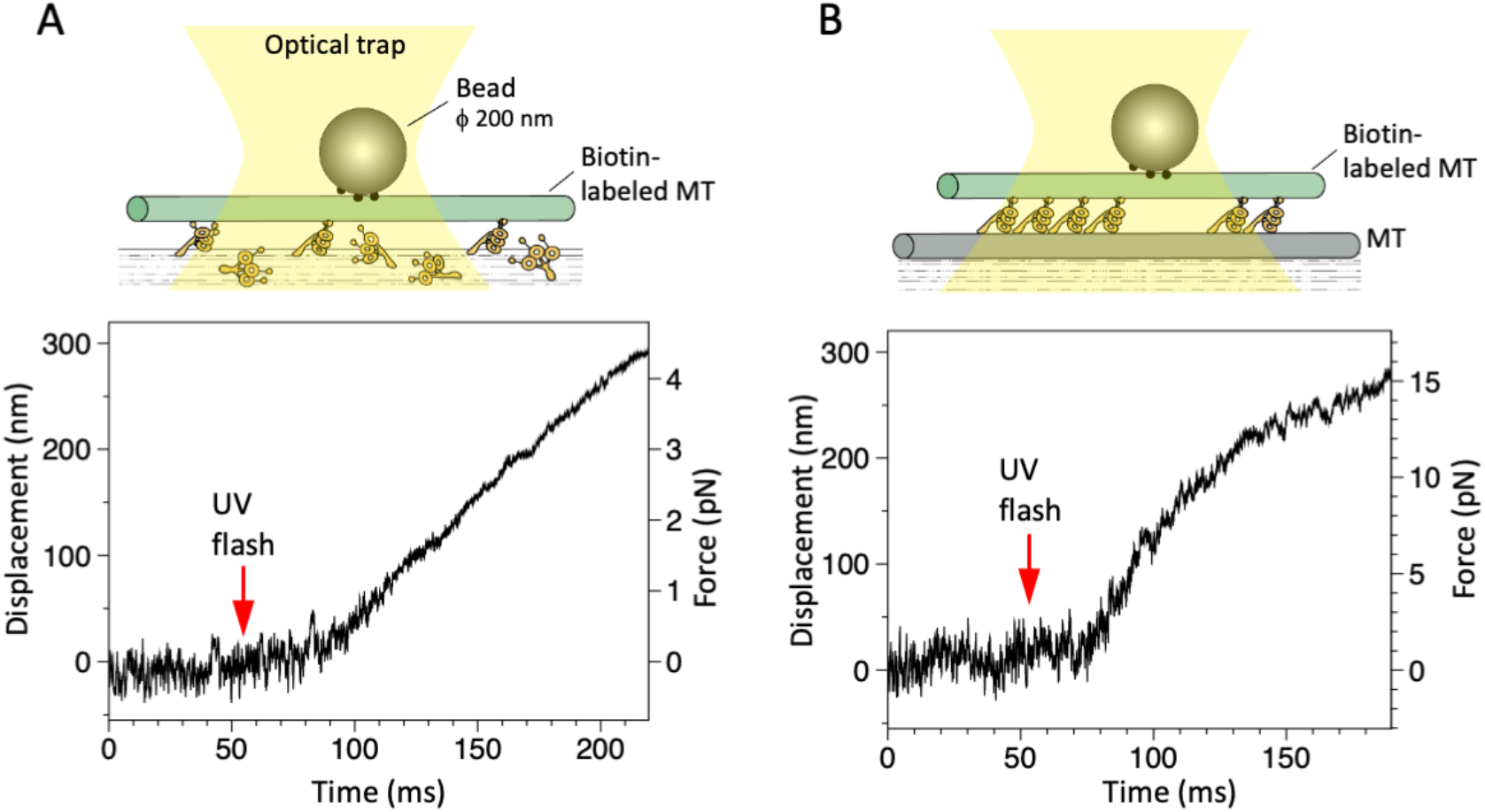
Motility in the absence of oppositely oriented dyneins measured in optical trapping assays. (A) Displacement of a MT in a MT-gliding assay. Trap stiffness: 0.015 pN/nm. (B) Displacement of a MT in the dynein-MT complex in which dyneins are arranged unidirectionally (see Fig S2C). Trap stiffness: 0.055 pN/nm.

For the experiments in Fig 5A, a bead (200 nm in diameter) was attached to a MT that was bound to the dynein-coated surface, and displacement after UV photolysis of caged ATP was measured in optical trapping assays. The number of dynein molecules interacting with a MT (4.7±1.9 µm (mean±SD)) was roughly estimated to be 24 at a dynein concentration of 12.5 µg/ml (see Materials and Methods). In contrast to the previous results using single or a few dyneins (Shingyoji *et al*., 1998; Shingyoji *et al*., 2015), the MTs moved smoothly in one direction (Fig 5A).

We have also examined the motility powered by a unidirectional array of dyneins. Dynein-MT-complexes that normally contain oppositely-oriented dyneins were disassembled by ATP, and new MTs were added in the absence of ATP to make dynein-MT complexes in which all the dyneins are oriented in the same way (illustrated in Fig S2C). The newly-added MTs were biotinylated to enable binding of avidin-coated beads in optical trapping experiments, and were also more brightly labelled so that they could be identified in the dynein-MT complex. In gliding assays the MTs in these complexes moved in one direction with an average speed of 6.2 µm/s, which is faster than those in the usual surface-gliding assay (Fig S2C), as reported previously (Aoyama & Kamiya, 2010). Optical trapping experiments also showed at a higher resolution that the movement was unidirectional (Figs 5B and S6), clearly different from the traces observed with the complex containing oppositely oriented dyneins (Figs 3B and S5). Thus, we conclude that an ensemble of multiple uniformly oriented dynein molecules moves a MT unidirectionally, and that two groups of oppositely oriented dyneins are required for oscillation.

### Structural changes of dynein in the dynein-MT-DNA-origami complex

During oscillatory movement, the two groups of oppositely oriented dynein molecules are likely to have different conformations. Previous structural analysis of axonemes revealed that the heads of dynein move toward the MT minus end in the presence of ATP or ADP·vanadate, or in beating flagella (Burgess, 1995; Lin & Nicastro, 2018; Movassagh *et al*., 2010; Sale *et al*, 1985; Ueno *et al*, 2014), so that the length of the dynein molecule measured along a MT increases. Preliminary analysis of our negative-stain EM images showed that the length increases for both groups of oppositely oriented dyneins in the presence of ATP (p<0.00001, T-test), indicating that their structures are both different from the rigor structure (Fig S7). Differences in the average lengths of two oppositely oriented dyneins were not detected. However, the arrangement of the three heads seemed to be different between the two oppositely oriented dyneins in the presence of ATP (Fig S7C), whereas those in the absence of ATP looked similar to each other (Fig S7B). Although the different arrangements of the heads in the presence of ATP might correspond to the different structural states observed on the opposite sides of sea urchin flagella (Lin & Nicastro, 2018), further structural analysis would be required to reveal the structural changes in detail.

### Bending motions of the dynein-MT-DNA-origami complex

In cilia and flagella, which are free to move in water, relative movements of the neighboring doublet MTs lead to bending motions. On the other hand, the dynein-MT-DNA-origami complexes in our optical trapping experiments were fixed to the glass surface along the whole length. In order to investigate whether the dynein-MT-DNA-origami complex has an ability to produce bending motions, we searched for a complex which is held at one point. Whereas a majority of the complexes seemed to be fixed to the glass along their entire length, there were some complexes attached only partially either to a bead or to the glass. Some examples are shown in Figs 6 and S8, and Movies S3 and S4. Although our fluorescence microscopy setup did not allow us to detect high-frequency oscillations, repetitive bending motions were indeed observed. The plot in Fig 6B shows that the complex can bend in both directions compared to the position before ATP release. Thus, it is likely that the bending motions in the opposite directions are powered by oppositely oriented dyneins, not just the cycles of force production and detachment. Sometimes a part of the MT bundle separated during bending motions as in Fig S8A, probably because this region did not have DNA origami attachment or because the DNA origami was pulled away during bending. Using these regions, relative sliding distances of the MTs were estimated to be approximately 130 nm and 365 nm for the frames #38 of Fig 6A and #13 of Fig S8, respectively. Sliding distances longer than the expected maximum value (~200 nm) are thought to be caused by rearrangement of the DNA origami during MT sliding. The bending motions observed here indicate that a system composed of MTs, oppositely oriented dyneins, and inter-MT crosslinkers has the ability to bend repetitively.

**Figure 6.**
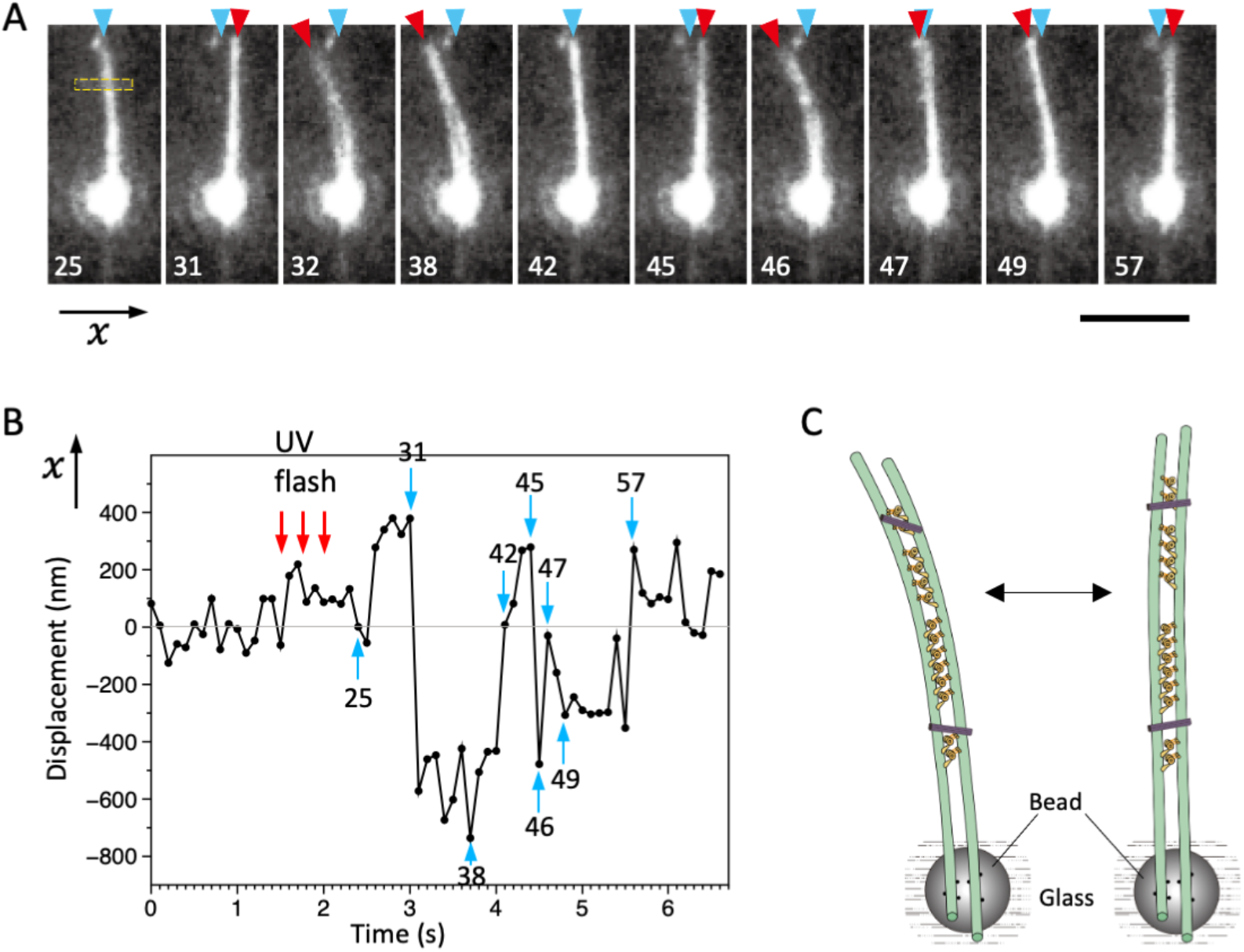
Bending motions of the dynein-MT-DNA-origami complex. (A) Snapshots during movement of the dynein-MT-DNA-origami complex shown in Movie S3. The number indicated in each frame corresponds to the frame number in the movie (recorded at 10 frames/s). The UV was flashed at frames #16, 18, and 21, each for 50 ms. The complex is attached to a bead while one end (red arrowheads) is free, which moves with respect to the position before the UV flash (cyan arrowhead) as the complex bends repeatedly. Bar, 5 µm. (B) Displacement of the complex during bending motion. The plot shows the lateral (x) positions of the complex observed in the boxed region in A. The average position before the UV flash (frames #5 - 15) was taken as 0 on the vertical axis. Note that the displacement is to both the plus and minus directions. (C) Diagram illustrating a model explaining the movement.

## Discussion

Repetitive bending of cilia and flagella is produced by collective functions of many dynein molecules that are aligned in a specific manner and communicate with each other. Another important feature of cilia and flagella is that the dyneins on the opposite sides of the axoneme produce force to bend the axoneme in opposite directions. The dynein-MT system used here mimics cilia and flagella in that it contains two groups of oppositely-oriented dynein molecules, that the neighboring dyneins are arranged with 24-nm periodicities between a pair of MTs, and that the MTs are interconnected with passive linkers. Previous work using frayed *Chlamydomonas* flagella axonemes showed that a pair of doublet MTs can display cyclic association, buckling, and dissociation (Aoyama & Kamiya, 2005; Brokaw, 2009). In their system, the doublet MTs were bundled at the proximal end and the inter-doublet linkers were proteolyzed. Our dynein-MT complex differs from theirs in that it contains inter-MT linkers and two groups of dyneins that produce opposing forces between a pair of MTs.

Oscillatory movements of MTs were also observed by Shingyoji et al.(Shingyoji *et al*., 1998; Shingyoji *et al*., 2015), using single or a few molecules of sea urchin axonemal dynein. In contrast, our work shows that multiple molecules of *Chlamydomonas* outer-arm dynein move a MT unidirectionally. We think the discrepancy is caused by the number and the arrangement of molecules: our system has multiple dynein molecules (~35 on average in the case of the complex with oppositely oriented dyneins) aligned regularly on a MT, which would work cooperatively as in cilia and flagella. Effect of the number of molecules on motor’s directionality was also reported for kinesin-5 Cin8: a single Cin8 moved to the MT minus end, while a team of Cin8 showed plus-end-directed motility (Roostalu *et al*, 2011). Our results show that an ensemble of dynein molecules aligned on a MT produce force in one direction, but participation of oppositely oriented dyneins is required for oscillatory movements.

The axoneme is a complex structure containing various components that may modulate the dynein activity, which makes it difficult to understand the basic mechanism of oscillation. For example, radial spokes and central apparatus are thought to be important for beating of flagella, but mutant flagella lacking radial spokes or the central pair can beat under certain conditions (Frey *et al*, 1997; Yagi & Kamiya, 2000), indicating that they are not absolutely needed for oscillatory beating. The circumferential layout of nine doublet MTs and/or the cooperative activities of outer-arm and several species of inner-arm dyneins may also be important for oscillation, but it has not been known whether they are essential for oscillation. Some modeling studies predicted that a system composed of dynein, MTs, and passive components can produce oscillation due to mechanical instability (Bayly & Dutcher, 2016), but direct evidence has not been obtained because of the lack of the reconstituted system with minimal components. Here, use of DNA origami enabled us to construct a simple system composed of two MTs with clusters of oppositely oriented dyneins and passive linkers. Oscillation of this system show that the regulatory components including the 9+2 structure with radial spokes and the central apparatus, nexin-dynein regulatory complexes, and inner-arm dyneins are not essential for generation of oscillation.

Since the dynein-MT complex with unidirectionally aligned dynein did not show oscillation, the most likely explanation for the oscillatory movements would be that each of the two dynein groups are responsible for forward and backward movements, respectively (Fig 1D). The DNA linkers limit the range of relative sliding, and thus may function as the trigger to change the direction of movement. The average frequency of the observed oscillation (~33 Hz) was close to the beat frequencies of the two flagella of a detergent-extracted *Chlamydomonas* cell (30 and 45 Hz) (Kamiya & Hasegawa, 1987). In addition, the dynein-MT-DNA-origami complex showed repetitive bending motions when one end is free in solution. Although regulatory components including the radial spokes and central apparatus are probably needed for more coordinated and long-lasting oscillation, our results show that a minimal system composed of two groups of oppositely oriented dyneins, MTs, and inter-MT crosslinkers has an intrinsic ability to cause oscillation and cyclic bending motions.

The dynein-MT-DNA-origami complex developed here contains multiple dynein molecules aligned between the MTs with a 24-nm periodicity as in axonemes, allowing the neighboring dyneins to cooperate, and the oppositely oriented dyneins to work against each other. This simple system is thus useful for the future research of the mechanism of cilia/flagella motility. It can also be used to study the high-resolution structures of dynein during oscillatory movement. Preliminary results of negative-stain EM analysis suggested that the dynein molecules oppositely oriented between a pair of MTs are in different structural states. Future cryo-EM studies analyzing the 3D structures of activated and regulated dynein molecules would be of great interest.

## Materials and Methods

### DNA origami

Single-stranded P8064 DNA purchased from *tilibit nanosystems GmbH* (*Garching, Germany)* was used as the scaffold for DNA origami. Unmodified staple strands were purchased from Sigma-Genosys as Oligonucleotide Purification Cartridge (OPC) grade. Amino-modified staples, which have three amino-modified C6dT oligonucleotide near the 5’ end, were purchased from IDT as Dual-HPLC-purified, or from Japan Bio Service (Tsukuba, Japan) as HPLC-purified. The SNAP-ligand was covalently attached to the amino-modified staples by mixing the staples and BG-GLA-NHS (NEB, dissolved in DMSO) as previously described (Derr *et al*, 2012; Masubuchi *et al*, 2018). The label efficiency of the SNAP-ligand was estimated to be 95−98% by a gel-shift assay using purified SNAP-tag protein. The modified staples are described in Table S1.

The rod type DNA origami nanostructure composed of 30 helices was designed using the honeycomb-lattice version (10.5 bp/turn) of the caDNAno2 software, and folded in 1 × Rod buffer (5 mM Tris boric acid, pH 7.6, 20 mM Mg(OAc)_2_, 5 mM Na(OAc), and 1 mM EDTA). Typically, 40 nM single-stranded P8064 DNA (tilibit) and 240 nM of each staple strand (6-fold excess) were mixed in 1 × Rod buffer, denatured at 85°C for 5 minutes and annealed at 47.5°C for 4 hours in a PCR machine (Takara-Bio). The folded DNA origami was then agarose-gel-purified (Douglas *et al*, 2009), and concentrated by PEG-precipitation (Stahl *et al*, 2014). The Rod concentration was estimated by absorbance at 280 nm using NanoDrop (Thermo).

### Preparation of the Dynein-MT-DNA-origami complex

Dynein was extracted from *Chlamydomonas* flagella with high salt. Microtubules were polymerized using tubulin purified from brain tissue and taxol-stabilized. Detailed methods for preparation of dynein (Yagi *et al*, 2009), MTs, and kinesin (Miyazono *et al*, 2010) are described in Supplementary Method. 15-20 nM DNA rods were incubated with kinesin E237A at a molar ratio of 1:8 in HEM buffer (20mM HEPES, 1mM EGTA, 10mM MgSO_4_, pH= 7.8) for 30 minutes at room temperature. Meanwhile, the dynein-MT complex was prepared by incubating 10-20 µg/ml dynein preparation with ~40 µg/ml MTs in HEM for 8-10 minutes at room temperature. Dynein-MT and DNA-kinesin were then mixed so that the final concentrations of dynein, MTs, DNA, and kinesin were 6.25 or 12.5 µg/ml, 25 µg/ml, 2.5 nM, and 20 nM, respectively, and incubated for 10 minutes at room temperature. For the dynein-MT complex without DNA-kinesin, HEM was added instead of DNA-kinesin.

### Preparation of the Dynein-MT-DNA complex with a unidirectional array of dynein

For preparation of the dynein-MT complex that has dyneins oriented in the same way (Figs 5B and S2C), dynein-MT complexes were prepared using less-brightly-fluorescent MTs (rhodamine tubulin: unlabeled tubulin = 3:97) at a final concentration of dynein and MTs of 30 µg/ml and 15 µg/ml, respectively, and adsorbed to the glass surface of a chamber. The complexes, which are thought to contain oppositely oriented dyneins, were then separated by addition of 0.5 mM ATP in HEM. After perfusion of HEM containing 1 mM ADP, brighter MTs (rhodamine tubulin: biotinylated tubulin: unlabeled tubulin = 1:2:7) were added to make dynein-MT complexes that contain dyneins in a single orientation. Biotinylated tubulin was used for binding to avidin-coated beads in optical trapping experiments.

### EM and image analysis

The sample was applied onto a carbon-coated grid and stained with 1-2 % uranyl acetate. For observing the complex in the presence of ATP, ADP (final: 1 mM) was first added to the sample on a grid (Yagi, 2000). After 1 min, ATP (final: 0.1 mM) was added and the sample was immediately stained with uranyl acetate. The samples were observed using an FEI Tecnai F-20 electron microscope equipped with a Gatan Orius 831 CCD camera. The images were adjusted for contrast and Gaussian-filtered in Adobe Photoshop to reduce noise. The images of individual dynein molecules were extracted and analyzed using the Eos software (Yasunaga & Wakabayashi, 1996), as described in Supplementary Method.

### Motility assays and optical trapping

The detailed methods of preparation of the glass chambers and MT-gliding assays are described in Supplementary Method. Optical trapping experiments were performed as previously described using a custom-made microscope system (Kinoshita *et al*, 2018). After perfusion of the dynein MT-complex with and without DNA-kinesin, 1mM ADP in the HEM buffer, and then the assay buffer (1 mM caged ATP (Dojin), 0.1 mM taxol, 20 mM glucose, 0.5 % (v/v) β-mercaptoethanol, 20 mg/ml catalase, 100 μg/ml glucose oxidase, and 1 unit/ml hexokinase in HEM) with streptavidin-coated beads and 0.2 mg/ml casein were introduced into the glass chamber.

A bead was trapped and then brought in contact with a MT in the complex. Caged ATP was photolyzed by a UV laser pulse. Displacement of the bead was measured at a sampling rate of 20 kHz with a quadrant photodiode connected to a MacLab system (AD Instruments). For the experiments with dynein-MT-DNA-origami complexes, 200 nm beads were used (trap stiffness: 0.015-0.055 pN/nm). For measuring the maximum force produced by the dynein array in the dynein-MT complex (Fig S3), 500 nm beads were used (trap stiffness: 0.175-0.35 pN/nm), because the bead-attached MT often slid out of the optical trap and the dynein-MT complex disassembled when 200 nm beads were used.

For the optical trapping experiments of MTs gliding over dynein-coated surfaces (Fig 5A), 10 µl of 125 µg/ml dynein was applied to the glass chamber as in usual MT-gliding assays. Assuming that 10 % of the perfused dynein molecules are adsorbed to the glass (Sakakibara *et al*., 1999), the density of the dynein molecules was roughly estimated to be 209 molecules/µm^2^. Thus, if dynein molecules directly below a MT that has a diameter of 25 nm can interact with the MT, the number of dynein molecules that interact with the MT is calculated to be 5.2 molecules/µm.

The steps were detected by a step-finding algorithm by Kerssemakers et al. (Kerssemakers *et al*., 2006), as described in Supplementary Method. Fitting with Gaussian functions was performed by graphic software (Datagraph, Visual Data Tools, Inc.).

### Observation of bending motions

Avidin-coated beads (200 nm in diameter) were perfused into a glass chamber. After washing with HEM buffer, dynein-MT-DNA-origami complexes prepared as above using 6.25 µg/ml (final) dynein and rhodamine-labeled MTs were introduced into a chamber and incubated for 5 min. The assay buffer containing 1 mM caged ATP, 0.5 mM ADP, and 0.3 mg/ml casein was then perfused and the sample was observed under a fluorescence microscope. Although the majority of complexes firmly attached to the glass and did not show movement, some complexes appeared to be attached only by a part of their length while the other ends were free and showed movement. For some experiments, avidin-coated beads were first introduced to the chamber, and dynein-MT-DNA-origami complexes made with rhodamine and biotin-labeled MTs were bound to the beads.

Positions of the MTs were detected by fitting Gaussian functions to the fluorescence intensity profile using ImageJ. The maximum values of relative sliding of the MTs in the complex were estimated using the parts where 2 MTs of the complex separated (e.g., shown by orange arrowheads in Fig S8A). The length along each MT was measured and the difference between the lengths of the two MTs was taken as the sliding distance.

## Acknowledgments

We are grateful to T. Yagi, K. Wakabayashi and R. Kamiya for the gift of *Chlamydomonas* strains and continuous advice on dynein preparation, C. Shingyoji and I. Nakano for guidance on motility assays, A. Nagasaki for advice on fluorescence microscopy, Y. Harada for support, and H. Ueno for initial experiments with the dynein-MT system. This work was supported by JSPS KAKENHI Grant Numbers 24115522, 16K07332, 17H05898 for KH, 15H00798, 19H03197 for HT, 16H04773, 19H03189 for HH.

## Author contributions

K.H. designed the outline of the study. S.A.A., K.Y., and K.H. prepared the dynein-MT complexes. H.Tad. and K.F. designed and prepared the DNA origami linkers. S.A.A., H.H. and Y.K. carried out motility and force measurements. S.A.A., T.F., Y.K., H.H., and K.H. analyzed the motility data. K.H, Y.K, and R.B. carried out electron microscopy. H.Tak., T.Y., and K.H. analyzed the electron micrographs. K.H., S.A.A., H.Tad., and H.H. discussed the data and wrote the manuscript.

## References

Alper JB, Tovar M, Howard J (2013) Displacement-Weighted Velocity Analysis of Gliding Assays Reveals that Chlamydomonas Axonemal Dynein Preferentially Moves Conspecific Microtubules. Biophys J 104: 1989–1998

Aoyama S, Kamiya R (2005) Cyclical interactions between two outer doublet microtubules in split flagellar axonemes. Biophys J 89: 3261–3268

Aoyama S, Kamiya R (2010) Strikingly fast microtubule sliding in bundles formed by Chlamydomonas axonemal dynein. Cytoskeleton (Hoboken) 67: 365–372

Bayly PV, Dutcher SK (2016) Steady dynein forces induce flutter instability and propagating waves in mathematical models of flagella. J R Soc Interface 13

Belyy V, Hendel NL, Chien A, Yildiz A (2014) Cytoplasmic dynein transports cargos via load-sharing between the heads. Nat Commun 5: 5544

Brokaw CJ (2009) Simulation of Cyclic Dynein-Driven Sliding, Splitting, and Reassociation in an Outer Doublet Pair. Biophys J 97: 2939–2947

Burgess SA (1995) Rigor and relaxed outer dynein arms in replicas of cryofixed motile flagella. J Mol Biol 250: 52–63

Camalet S, Jülicher F (2000) Generic aspects of axonemal beating. New J Phys 2: 24.21-24.23

Derr ND, Goodman BS, Jungmann R, Leschziner AE, Shih WM, Reck-Peterson SL (2012) Tug-of-war in motor protein ensembles revealed with a programmable DNA origami scaffold. Science 338: 662–665

Douglas SM, Dietz H, Liedl T, Hogberg B, Graf F, Shih WM (2009) Self-assembly of DNA into nanoscale three-dimensional shapes. Nature 459: 414–418

Frey E, Brokaw CJ, Omoto CK (1997) Reactivation at low ATP distinguishes among classes of paralyzed flagella mutants. Cell Motil Cytoskeleton 38: 91–99

Furuta A, Yagi T, Yanagisawa HA, Higuchi H, Kamiya R (2009) Systematic comparison of in vitro motile properties between Chlamydomonas wild-type and mutant outer arm dyneins each lacking one of the three heavy chains. J Biol Chem 284: 5927–5935

Furuta K, Furuta A, Toyoshima YY, Amino M, Oiwa K, Kojima H (2013) Measuring collective transport by defined numbers of processive and nonprocessive kinesin motors. Proc Natl Acad Sci U S A 110: 501–506

Gennerich A, Carter AP, Reck-Peterson SL, Vale RD (2007) Force-induced bidirectional stepping of cytoplasmic dynein. Cell 131: 952–965

Goodenough UW, Heuser JE (1982) Substructure of the outer dynein arm. J Cell Biol 95: 798–815

Haimo LT, Telzer BR, Rosenbaum JL (1979) Dynein binds to and crossbridges cytoplasmic microtubules. Proc Natl Acad Sci U S A 76: 5759–5763

Heuser T, Raytchev M, Krell J, Porter ME, Nicastro D (2009) The dynein regulatory complex is the nexin link and a major regulatory node in cilia and flagella. J Cell Biol 187: 921–933

Hirakawa E, Higuchi H, Toyoshima YY (2000) Processive movement of single 22S dynein molecules occurs only at low ATP concentrations. Proc Natl Acad Sci U S A 97: 2533–2537

Hirose K (2012) Structural Analysis of Dynein Bound to Microtubules. Handbook of Dynein: 81–97

Kamiya R, Hasegawa E (1987) Intrinsic difference in beat frequency between the two flagella of Chlamydomonas reinhardtii. Exp Cell Res 173: 299–304

Kerssemakers JWJ, Munteanu EL, Laan L, Noetzel TL, Janson ME, Dogterom M (2006) Assembly dynamics of microtubules at molecular resolution. Nature 442: 709–712

Kinoshita Y, Kambara T, Nishikawa K, Kaya M, Higuchi H (2018) Step sizes and rate constants of single-headed cytoplasmic dynein measured with optical tweezers. Sci Rep 8

Lin J, Nicastro D (2018) Asymmetric distribution and spatial switching of dynein activity generates ciliary motility. Science 360: eaar1968

Masubuchi T, Endo M, Iizuka R, Iguchi A, Yoon DH, Sekiguchi T, Qi H, Iinuma R, Miyazono Y, Shoji S et al (2018) Construction of integrated gene logic-chip. Nature Nanotechnology 13: 933–940

Mitchison TJ, Mitchison HM (2010) Cell biology: How cilia beat. Nature 463: 308–309

Miyazono Y, Hayashi M, Karagiannis P, Harada Y, Tadakuma H (2010) Strain through the neck linker ensures processive runs: a DNA-kinesin hybrid nanomachine study. EMBO J 29: 93–106

Movassagh T, Bui KH, Sakakibara H, Oiwa K, Ishikawa T (2010) Nucleotide-induced global conformational changes of flagellar dynein arms revealed by in situ analysis. Nat Struct Mol Biol 17: 761–767

Oda T, Hirokawa N, Kikkawa M (2007) Three-dimensional structures of the flagellar dynein-microtubule complex by cryoelectron microscopy. J Cell Biol 177: 243–252

Reck-Peterson SL, Yildiz A, Carter AP, Gennerich A, Zhang N, Vale RD (2006) Single-molecule analysis of dynein processivity and stepping behavior. Cell 126: 335–348

Rice S, Lin AW, Safer D, Hart CL, Naber N, Carragher BO, Cain SM, Pechatnikova E, Wilson-Kubalek EM, Whittaker M et al (1999) A structural change in the kinesin motor protein that drives motility. Nature 402: 778–784

Riedel-Kruse IH, Hilfinger A, Howard J, Julicher F (2007) How molecular motors shape the flagellar beat. HFSP J 1: 192–208

Roostalu J, Hentrich C, Bieling P, Telley IA, Schiebel E, Surrey T (2011) Directional switching of the kinesin Cin8 through motor coupling. Science 332: 94–99

Sakakibara H, Kojima H, Sakai Y, Katayama E, Oiwa K (1999) Inner-arm dynein c of Chlamydomonas flagella is a single-headed processive motor. Nature 400: 586–590

Sale WS, Goodenough UW, Heuser JE (1985) The substructure of isolated and in situ outer dynein arms of sea urchin sperm flagella. J Cell Biol 101: 1400–1412

Shingyoji C, Higuchi H, Yoshimura M, Katayama E, Yanagida T (1998) Dynein arms are oscillating force generators. Nature 393: 711–714

Shingyoji C, Nakano I, Inoue Y, Higuchi H (2015) Dynein arms are strain-dependent direction-switching force generators. Cytoskeleton (Hoboken) 72: 388–401

Soppina V, Rai AK, Ramaiya AJ, Barak P, Mallik R (2009) Tug-of-war between dissimilar teams of microtubule motors regulates transport and fission of endosomes. Proc Natl Acad Sci U S A 106: 19381–19386

Stahl E, Martin TG, Praetorius F, Dietz H (2014) Facile and Scalable Preparation of Pure and Dense DNA Origami Solutions. Angew Chem Int Edit 53: 12735–12740

Toba S, Watanabe TM, Yamaguchi-Okimoto L, Toyoshima YY, Higuchi H (2006) Overlapping hand-over-hand mechanism of single molecular motility of cytoplasmic dynein. Proc Natl Acad Sci U S A 103: 5741–5745

Ueno H, Bui KH, Ishikawa T, Imai Y, Yamaguchi T, Ishikawa T (2014) Structure of dimeric axonemal dynein in cilia suggests an alternative mechanism of force generation. Cytoskeleton (Hoboken) 71: 412–422

Ueno H, Yasunaga T, Shingyoji C, Hirose K (2008) Dynein pulls microtubules without rotating its stalk. Proc Natl Acad Sci U S A 105: 19702–19707

Witman GB, Plummer J, Sander G (1978) Chlamydomonas Flagellar Mutants Lacking Radial Spokes and Central Tubules - Structure, Composition, and Function of Specific Axonemal Components. J Cell Biol 76: 729–747

Yagi T (2000) ADP-dependent microtubule translocation by flagellar inner-arm dyneins. Cell Struct Funct 25: 263–267

Yagi T, Kamiya R (2000) Vigorous beating of Chlamydomonas axonemes lacking central pair/radial spoke structures in the presence of salts and organic compounds. Cell Motil Cytoskeleton 46: 190–199

Yagi T, Uematsu K, Liu Z, Kamiya R (2009) Identification of dyneins that localize exclusively to the proximal portion of Chlamydomonas flagella. J Cell Sci 122: 1306–1314

Yasunaga T, Wakabayashi T (1996) Extensible and object-oriented system Eos supplies a new environment for image analysis of electron micrographs of macromolecules. J Struct Biol 116: 155–160

Yokota E, Mabuchi I (1994) C/A dynein isolated from sea urchin sperm flagellar axonemes. Enzymatic properties and interaction with microtubules. J Cell Sci 107 (Pt 2): 353–361

