## Supplementary information for "Oscillatory movement of a dynein-microtubule complex crosslinked with DNA origami"

**This PDF file includes:**

Supplementary Figures S1 to S8
Legends for Supplementary Movies S1 to S4
Supplementary Methods
Supplementary Table S1

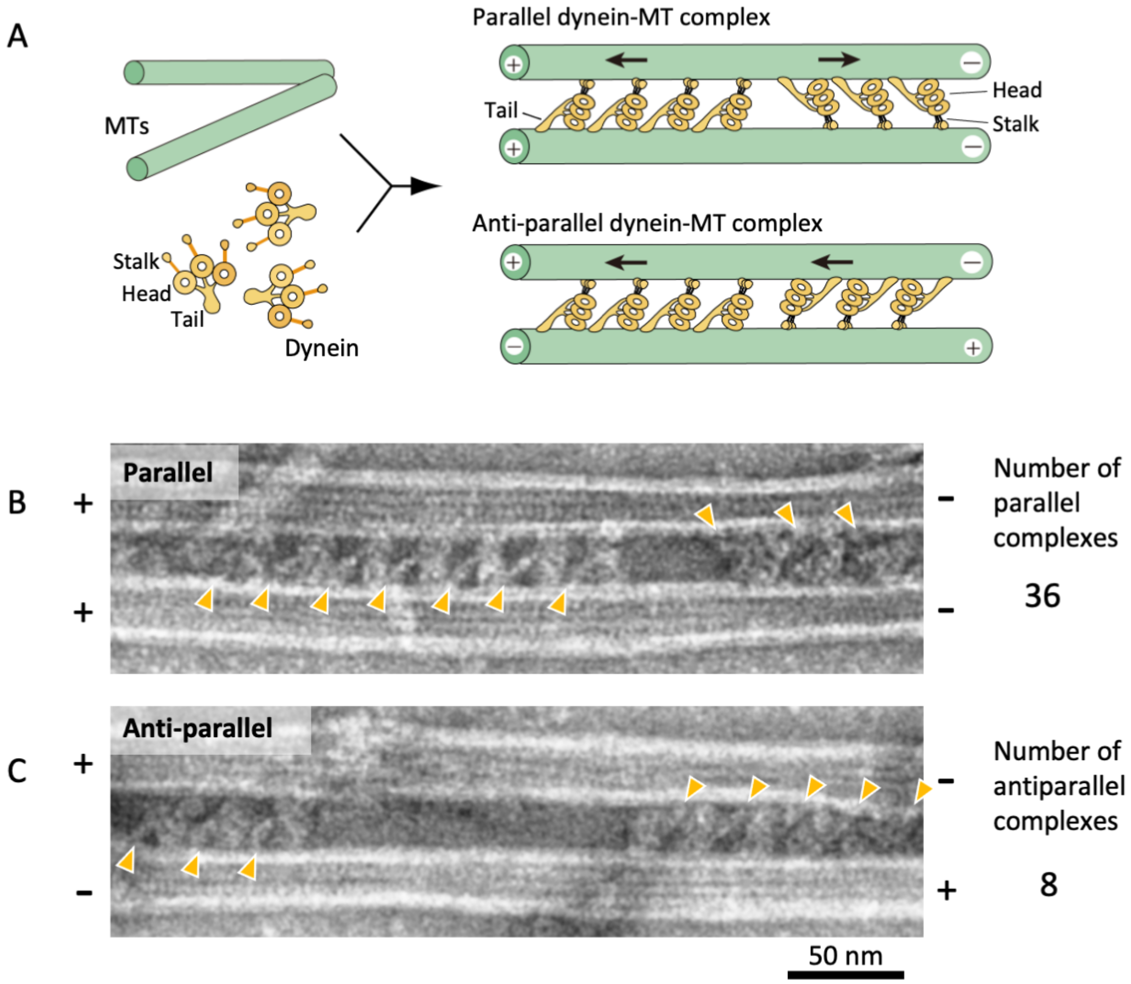


**Figure S1. Geometries of the dynein-MT complex.**

(A) A diagram showing two different geometries of the dynein-MT complex. An outer-arm dynein molecule cross-bridges two MTs by binding to one of the MTs with the MT-binding domain at the end of the stalk and to another MT with the tail. The stalk of each dynein is oriented closer to the minus end of the MT to which the stalk binds. Therefore, there are two possible orientations for dynein bound between a pair of MTs, depending on which MT its stalk binds to. If the two MTs have the same polarity as *in vivo*, two groups of dynein molecules produce opposing force (top-right). On the other hand, if the MTs are anti-parallel, all the dynein molecules produce force in the same direction (bottom-right).

(B and C) EM images of dynein-MT complexes showing two different geometries as illustrated in (A). When high-salt extracted *Chlamydomonas* dynein was used, ~80 % of the complexes had MTs with the same polarity, as judged by the negative stain images. The positions of the dynein tails are indicated by orange arrowheads.

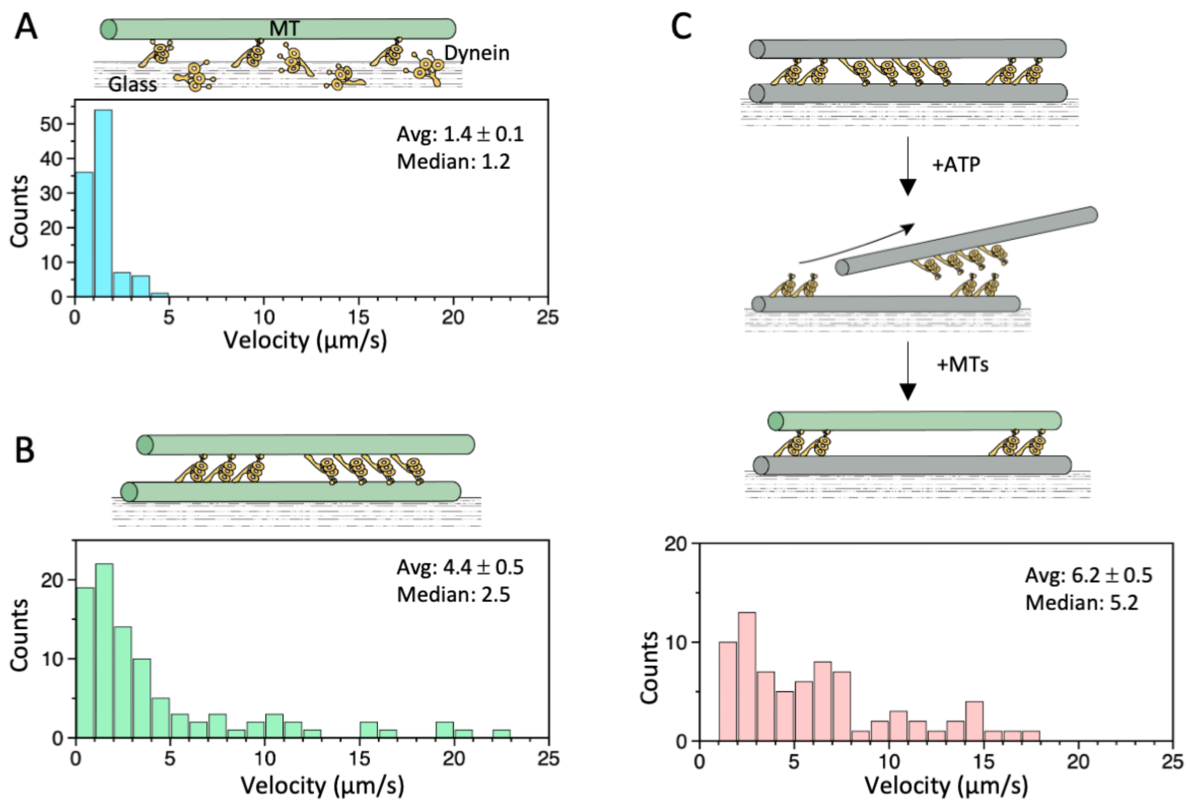


**Figure S2. MT-gliding velocities.**

Histograms of MT-gliding velocities in three different conditions. Average (mean ± s.e.m.) and median values of the velocities are indicated (N = 104, 94, and 74 for A, B, and C, respectively).

(A) Distributions of the MT gliding velocities in usual MT gliding assays in which MTs move over dynein-coated glass surfaces.

(B) Relative sliding of the MTs in a dynein-MT complex that contains dyneins in two opposite orientations.

(C) Relative sliding of the MTs in a dynein-MT complex in which dyneins are arranged unidirectionally. As illustrated in the diagram, dynein-MT complexes that contain dyneins in two opposite orientations were prepared using less brightly fluorescent MTs and adsorbed to the glass. Addition of ATP disassembles the complex, leaving MTs with dyneins oriented in the same way. New MTs (more brightly fluorescent) were added in the absence of ATP to make the complexes.

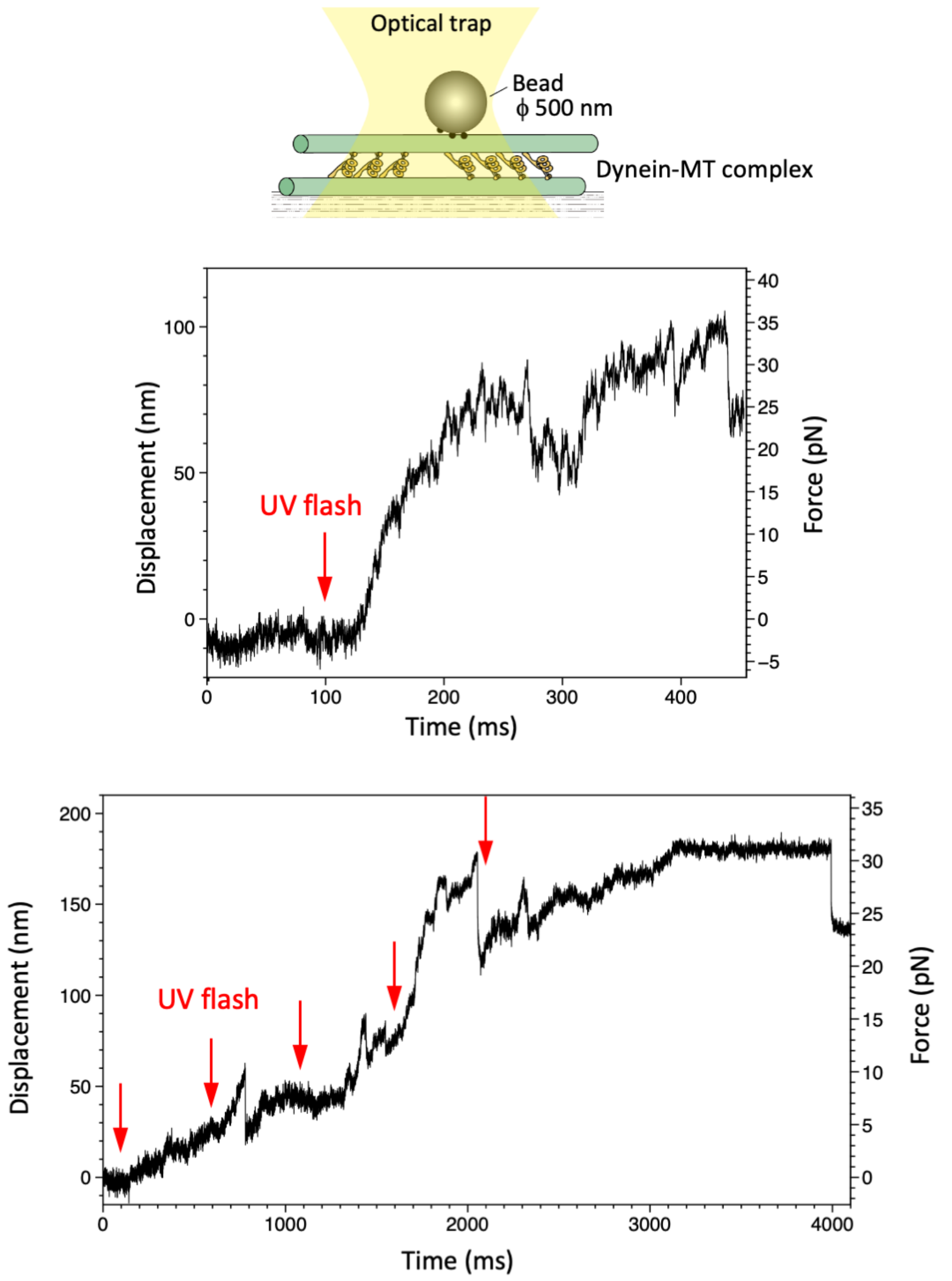


**Figure S3. Maximum force produced by the dynein-MT complex without DNA origami rods.**
Schematic representation of the experiment (not to scale) and two examples of the traces are shown. Larger beads (500 nm in diameter) were used to measure the maximum force. Red arrows indicate the timing of UV photolysis of caged ATP. In both cases, the maximum force was 30 – 35 pN.

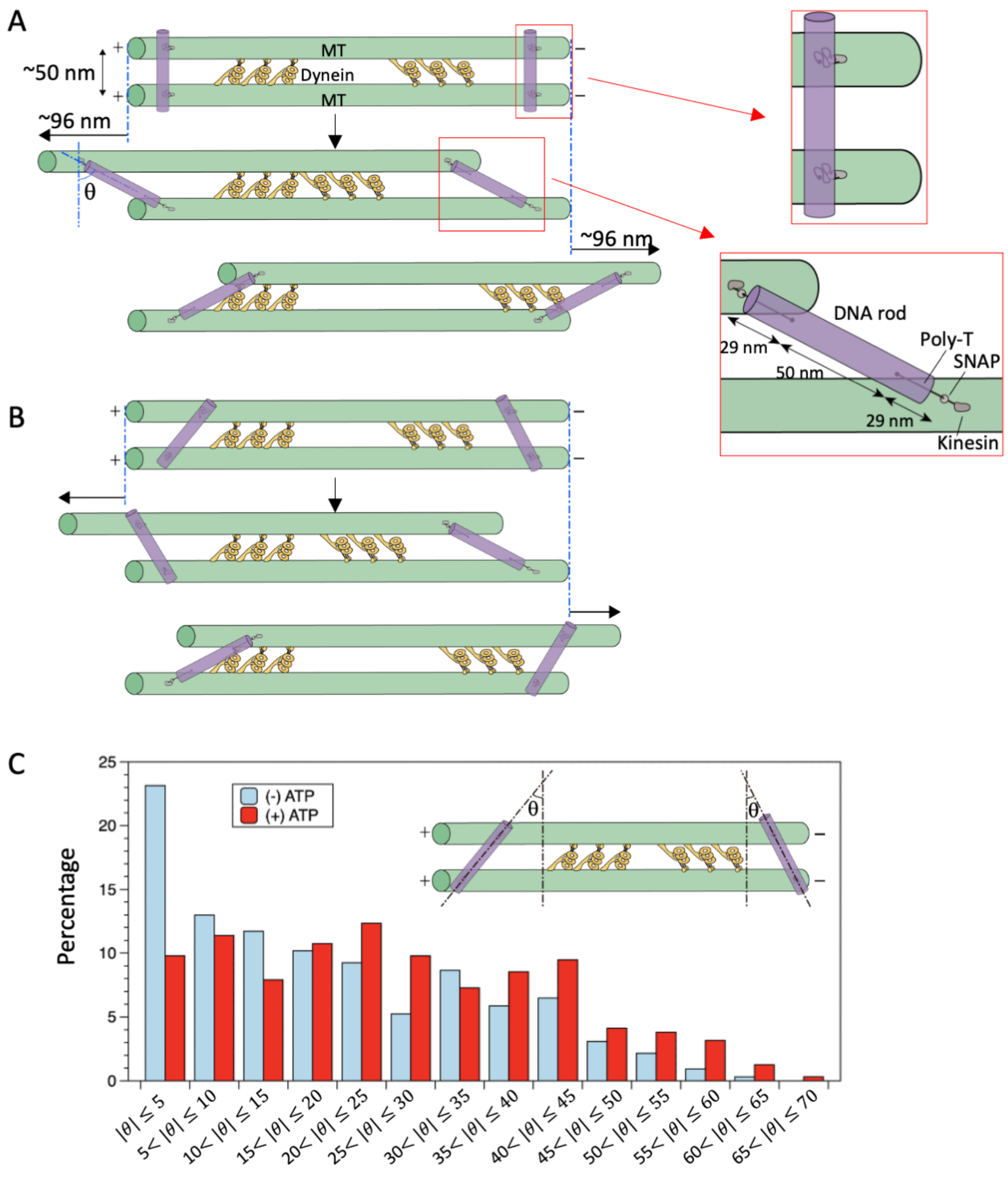


**Figure S4. Tilt angles of DNA origami rods.**

(A, B) When a DNA rod crosslinks two MTs, the initial binding angle is expected to have a distribution centered around 90 degrees with respect to the MTs. After addition of ATP, the MTs slide relative to each other, the linkers become stretched (illustrated in red boxes), and the DNA rods would tilt. Assuming the center-to-center distance between the MTs to be 50 nm and the length of the linker between DNA rods and the MT-binding site 29 nm (see Supplementary Methods), the maximum sliding distance and tilt angle ($\theta$) are calculated to be ~96 nm and ~62 degrees in both directions, when the initial tilt angle is 0 (A). However, when the initial binding angles are variable, the maximum sliding distance and tilt angle are expected to be smaller (B).

(C) Distributions of the tilt angles measured in the negative stain EM images (N=316, 324 for in the presence and absence of ATP, respectively). In the absence of ATP, the bin with tilt angle less than 5 degrees has the highest population. In the presence of ATP, the tilt angles have a wider distribution, with no apparent preference for smaller angles.

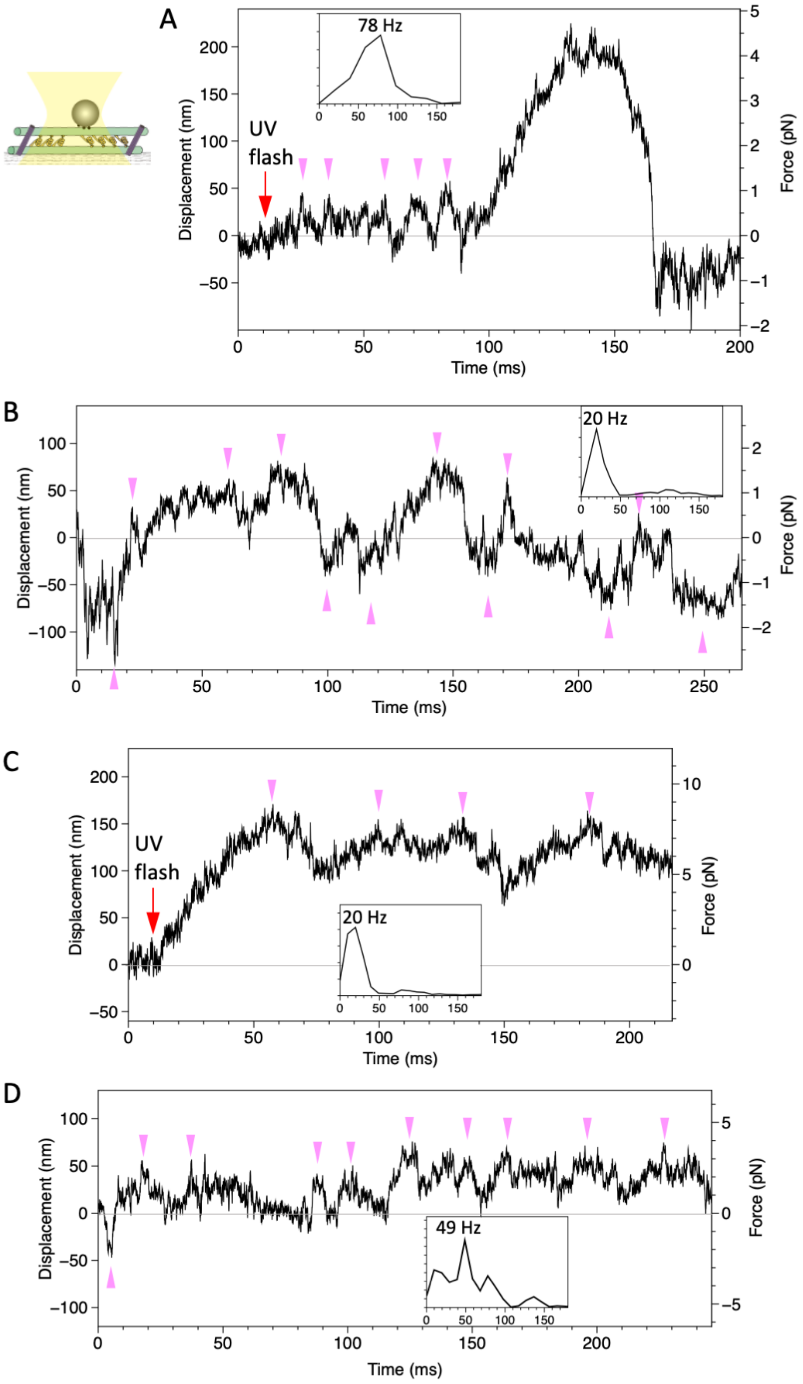


**Figure S5. Displacement of beads attached to the dynein-MT-DNA-origami complex.**

Four examples of traces, recorded under the similar conditions as in Fig 3 (trap stiffness 0.021 pN/nm for A and B, 0.052 pN/nm for C and D) are shown. In the trace in (A), the bead showed oscillatory movement (pink arrowheads) after photolysis of caged ATP and then moved for ~200 nm unidirectionally. The trace in (B) was recorded ~100 ms after the UV flash. The bead moved in both directions from the trap center (0 nm of the vertical axis). The insets show the power spectral density of the traces, with the peak value indicated.

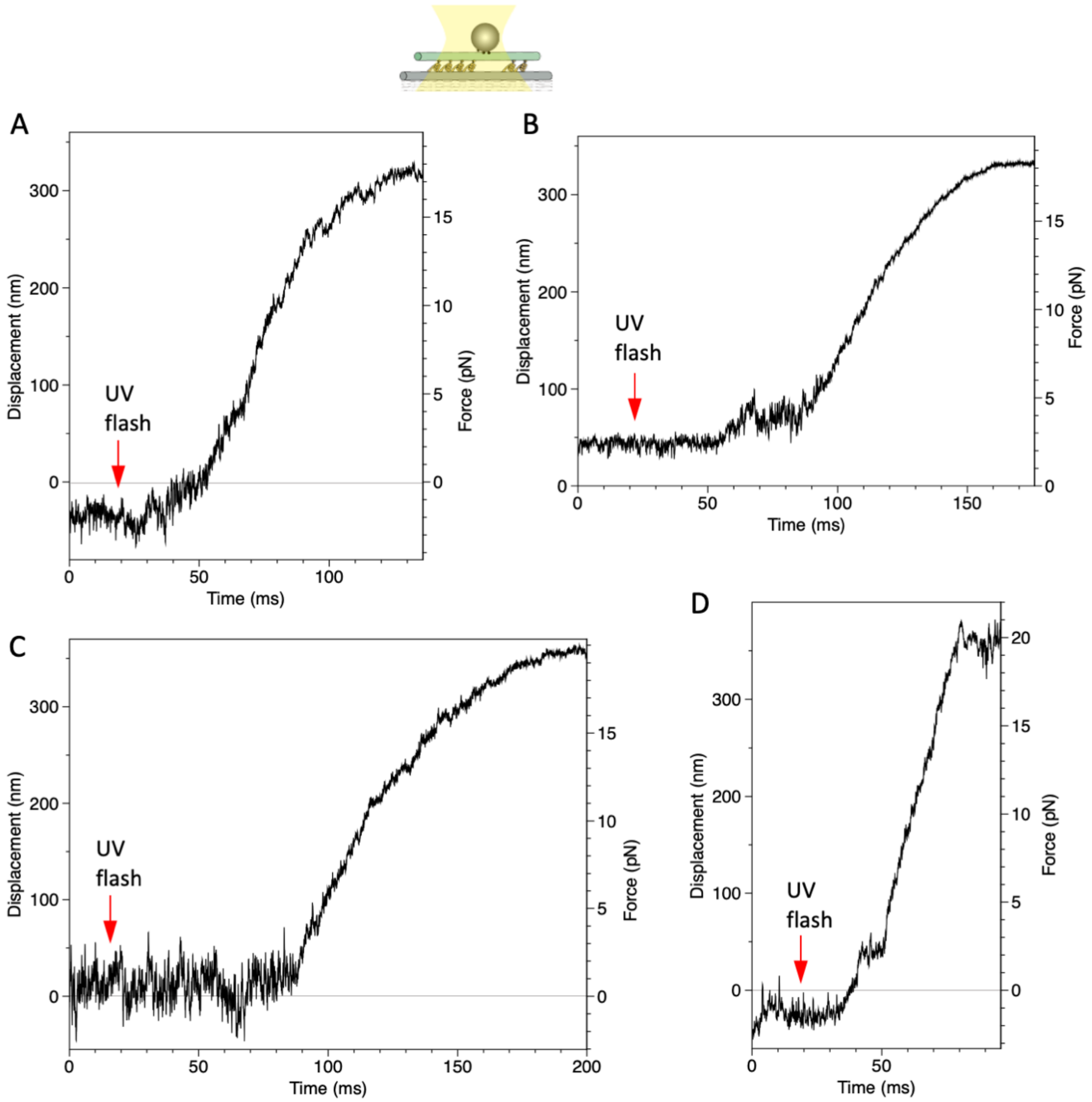


**Figure S6. Displacement of a bead attached to the dynein-MT complex in which dyneins are arranged unidirectionally.**

Four examples recorded under the same condition as in Fig 5B.

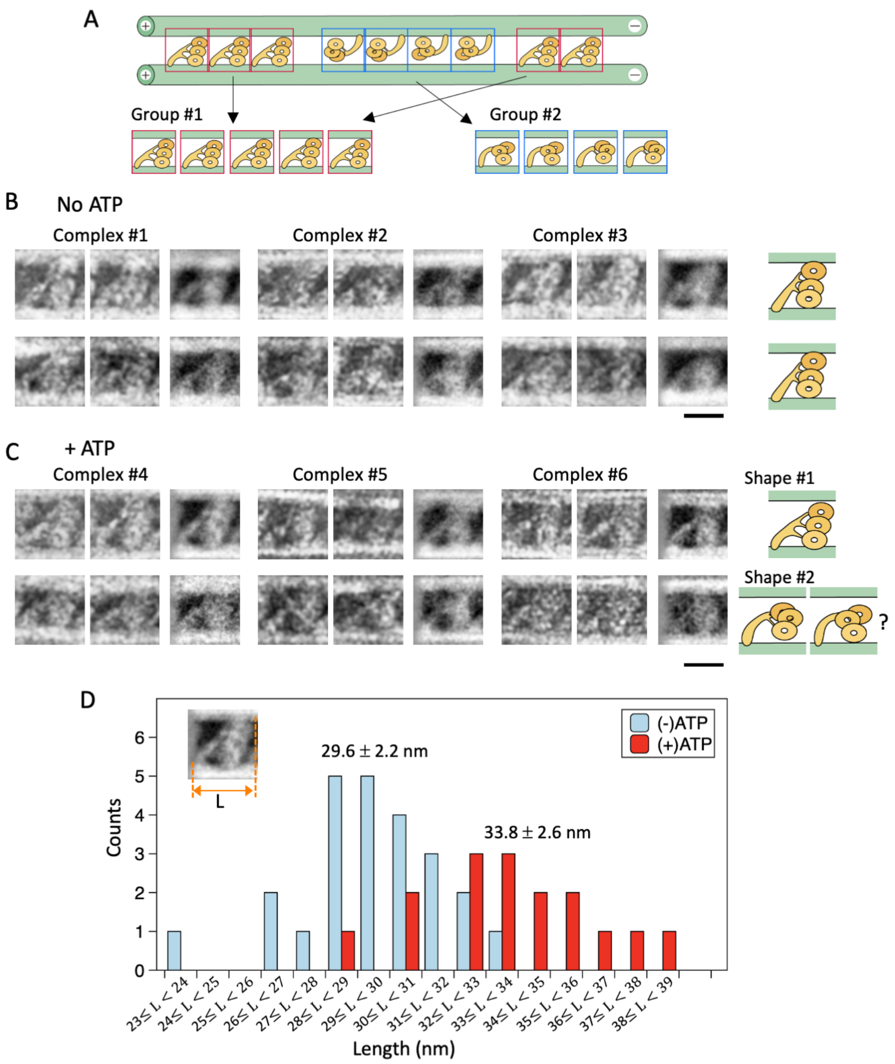


**Figure S7. Structural changes of dynein molecules in the absence and presence of ATP.**

(A) Schematic of the classification method. Images of individual dynein molecules in each dynein-MT-DNA-origami complex were classified into two groups depending on the orientation. The averaged image for each group was calculated.

(B, C) Examples of individual and averaged images. Images from three complexes in the absence of ATP (B) and three in the presence of ATP (C) are shown. The top and bottom rows show oppositely oriented dyneins, rotated so that the MT minus end is toward the right. For each group, two individual images (left) and the averaged image (right) are shown (N = 30 and 5, 11 and 11, 15 and 12, 22 and 2, 20 and 9, 14 and 19 for the complexes #1 to 6, respectively). A possible arrangement of the dynein heads is illustrated. Whereas the two averaged images look similar in the absence of ATP, those in the presence of ATP seem to have different arrangements of the dynein heads. The three heads in one of the groups are arranged in a straight line, while one of the heads in the other group appears to be shifted. Bar, 20 nm.

(D) Distribution of the averaged lengths (L) of MT-bound dynein molecules along the MT axis. Averaged lengths of the two groups of dyneins were calculated separately for each dynein-MT complex. The averaged length and standard deviation (n = 12 and 8 for the absence and presence of ATP, respectively) are indicated.

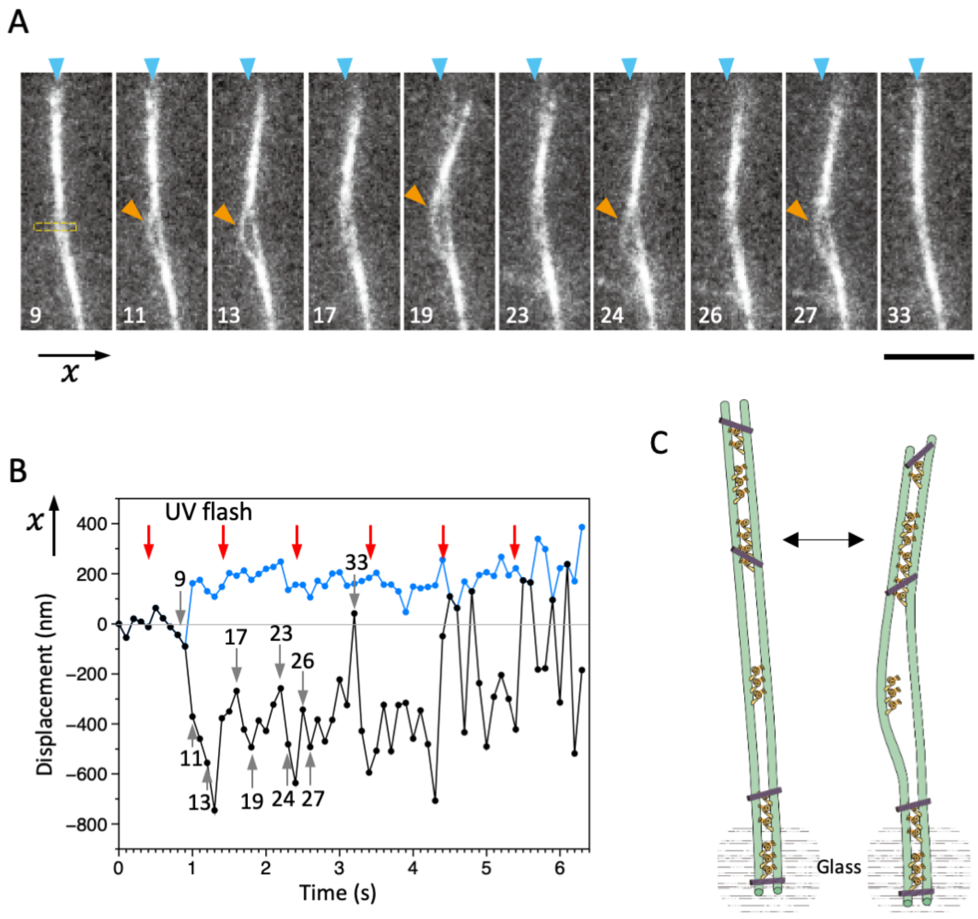


**Figure S8. Bending motions of the dynein-MT-DNA-origami complex**.

An example of the bending movement observed under the same experimental condition as in Fig 6 and Movie S3, but in this example, one end of the complex (bottom of the images) seemed to be attached to the glass. The movie of this complex is shown in Movie S4. The numbers indicated in A and B correspond to the frame numbers in Movie S4 (recorded at 10 frames/s).

(A) Snapshots during movement. UV was flashed at frames #5, 15, 25, 35, 45, and 55, each for 50 ms. After the UV flash, the middle part of the complex repetitively bows outward and becomes separated (orange arrowheads), presumably because there are not enough DNA origami and dynein molecules to hold the two MTs together when they are bent. Bar, 5 µm.

(B) Displacement of the complex during bending motion. The plot shows the lateral (x) positions of the two MTs (or MT bundles) observed in the boxed region in A, where the complex separated and one of the MTs bowed outward repeatedly. The average position before UV flash was taken as 0 on the vertical axis.

(C) Diagram illustrating a model explaining the movement.

### Legends for Expanded View Movies

**Movie S1.** Uni-directional movement of MTs from the dynein-MT complex. Dynein-MT complexes were adsorbed to the glass surface. Upon photolysis of caged ATP, some MTs (indicated by arrowheads) moved unidirectionally and slid out of the bundles. Recorded at 400 ms/frame.

**Movie S2.** Effect of DNA-origami rods on the movement of dynein-MT complexes. The dynein-MT-DNA-origami complex was adsorbed to the glass surface. The complex showed little movement after photolysis of caged ATP, indicating that relative sliding of MTs is restricted by cross-linking with DNA origami. Recorded at 400 ms/frame.

**Movie S3.** Bending motions of the dynein-MT-DNA-origami complex. Movie of the dynein-MT-DNA-origami shown in Fig 6. The complex is fixed to beads while one of the ends is free in solution. After UV photolysis of caged ATP (frames #16, 19, 21), the complex bends repeatedly. Recorded at 100 ms/frame.

**Movie S4.** Bending motions of the dynein-MT-DNA-origami complex. Movie of the dynein-MT-DNA-origami shown in Fig S8. Movement of the dynein-MT-DNA-origami complex was recorded in the same condition as in Movie S3 but without beads. One end of the complex is attached to the glass surface, while the MTs in the upper part of the image slide relative to each other, which causes bending and separation of the MTs in the middle part. UV flash at frames #5, 15, and 25. Recorded at 100 ms/frame.

### Supplementary Method

#### Preparation of dynein, kinesin, and microtubules

A wild-type strain (137c-) of *Chlamydomonas reinhardtii* was used. Flagella were isolated using dibucaine and demembranated as described previously (Yagi *et al*, 2009). For extraction of dynein, axonemes were incubated in 0.6 M KCl in HMDE solution (30 mM HEPES, 5 mM $MgSO_4$, 1 mM dithiothreitol, 1 mM EGTA, pH=7.4) for 30 minutes at 4°C. For some preparations, axonemes were first incubated in HMDEK (HMDE with 50 mM $CH_3COOK$) containing 0.1 mM ADP for 1 minute, centrifuged, and then incubated in 0.6 M KCl in HMDE for 20 minutes two times. After centrifugation, the excess salt was removed from the supernatant fraction either by overnight dialysis or by Amicon Ultra-4 100K centrifugal filter using HMDEK. The resulting dynein suspension was mixed with 23 % sucrose, aliquoted and stored in liquid nitrogen.

The E237A mutant of human ubiquitous kinesin, which is ATPase-defective with the neck-linker in a docked conformation (Rice *et al*, 1999), was used. SNAPf protein was fused to the C-terminus of cysteine-light, 336-residue monomer kinesin E237A. All constructs were verified by DNA sequencing. Monomeric kinesins were expressed and purified as previously described (Miyazono *et al*, 2010), except that kinesins were further purified with HiTrap-SP (GE) after His-tag purification (Ni-NTA Agarose, Qiagen).

Tubulin was purified from porcine brain tissue (Castoldi & Popova, 2003). Rhodamine-labeled and biotinylated tubulin was prepared using tetramethylrhodamine (C-1171, Molecular Probes) and Sulfo-NHS-LC-LC-Biotin (Thermo Scientific), respectively. For fluorescently-labeled MTs, rhodamine-labeled tubulin and unlabeled tubulin were mixed in the ratio 1:9. For MTs used to bind beads in the optical trapping experiments, rhodamine-labeled tubulin, biotinylated tubulin, and unlabeled tubulin were mixed in the ratio 1:2:7. MTs were polymerized in a polymerizing solution (80 mM PIPES (pH 6.8), 1 mM EGTA, 5 mM $MgSO_4$, 1 mM DTT, 0.5 mM GTP, and 5 or 10 % DMSO) for 30 min, stabilized with taxol and stored in liquid nitrogen.

#### Length estimation of the linker of the DNA origami

The maximum length of the linker was estimated assuming the following; the unit length of ssDNA : 0.63 nm, the unit length of amino acids : 0.34 nm, the unit length of a carbon chain : 0.13 nm, and the size of the SNAPf part : 2 nm. Using these values, the maximum length of the linker including 30 nucleotides of poly-thymidine, C6dT oligonucleotide, SNAPf, and an amino acid linker between kinesin and SNAPf, was calculated to be approximately 29 nm.

#### EM and image analysis

For the analysis of dynein shapes shown in Fig S7, a set of high magnification images and a low magnification image were taken from the same dynein-MT-DNA-origami complex. The high magnification images show the dynein structures at a higher magnification and comparison with the low magnification image tells where each dynein is located in the complex.

The electron microscopic images were analyzed using the Eos software (Yasunaga & Wakabayashi, 1996). Images of dynein-MT-DNA-origami complexes that have dynein molecules bound in two opposite orientations were selected. After correction for the contrast transfer function, image segments each containing a dynein molecule and parts of two MTs were extracted (Fig S7A). Images of dynein molecules whose tails are bound to the same MT were grouped, so that there are two groups of dynein images observed between a pair of MTs.

Image segments that belong to the same group were aligned and averaged in the following process. To lower the effect of the MTs and neighboring dyneins during alignment, the contrast of the peripheral region in each segment was reduced by 10%. Each segment was pasted into a slightly larger box filled with the mean density of the peripheral region. For each group, we selected one of the segments as the original reference and aligned the other images by calculating their similarity using a correlation function, allowing translation and rotation with 9 to 0.072-degree steps. The images in the same group were averaged using the rotation angles and translation values that showed the maximum correlation. We then used these average images as the references for the next cycle of alignment. The alignment cycle was repeated until the average images did not change.

The length of the dynein molecule along the MT axis was measured in each average image. The length of the dynein image projected onto the MT axis was defined as the dynein length (Fig S7D, inset). Distributions of the lengths in the presence and absence of ATP were compared statistically (Welch's T-test).

**Motility assays**

Glass chambers were made using two pieces of cover glass (24 mm x 32 mm (No. 24321, Muto pure chemicals, Japan) and 18 mm x 18 mm (No. 0101030, Marienfeld)). The glass was cleaned with KOH and then rinsed with $H_2O$ and ethanol for MT-gliding assays (Inoue & Shingyoji, 2007). For optical trapping experiments of the dynein-MT complex, un-cleaned glass was used because more beads adhered non-specifically to the cleaned glass. The size of the chamber was ~18 mm x 5 mm.

MT-gliding assays over dynein-coated glass surfaces (Fig S2A) were performed by introducing 100-125 μg/ml dynein in a chamber for 2 minutes, followed by 1 mM ADP in HEM for 1 minute, and then 5 μg/ml MT in the assay buffer (1 mM caged ATP (Dojin), 0.1 mM taxol, 20 mM glucose, 0.5 % (v/v) β-mercaptoethanol, 20 mg/ml catalase, 100 μg/ml glucose oxidase, and 1 unit/ml hexokinase in HEM). Movement of MTs after UV photolysis of caged ATP was observed using a fluorescence microscope (81X, Olympus, Japan) equipped with a 100X lens, a mercury lamp (USH1030L), and filters (535BP/30 and 330WB/80). Caged ATP was photolyzed by switching the filters. Images were recorded using a CMOS camera (ORCA-Flash 2.8, Hamamatsu Photonics, Japan).

For motility assays of the dynein-MT complex with and without DNA (Fig S2B and Movies S1 and S2), the complex preformed in solution was perfused into the chamber. After applying the ADP solution and assay buffer (without MTs), the MT movement after caged ATP photolysis was observed as above.

**Analysis of the displacement in optical trapping experiments**

Stepping events were analyzed by a step-finding algorithm by Kerssemakers et al. (Kerssemakers *et al*, 2006). To statistically optimize the step finding procedure, they defined a parameter *S*, which is the ratio of the $\chi^2$ of the counter fit to $\chi^2$ of the best fit. *S* changes with number of steps, and the highest peak S gives the best fit. For the analysis shown in Fig 4, only those data that had a clear peak in *S* were used. The detected step sizes were calibrated with attenuation factors to account for the compliance of the bead-MT linkage and that of dynein (Kinoshita *et al*, 2018). For calculation of the velocity during oscillation, we chose the peaks that showed at least 2 steps and amplitude ≥16 nm.

**References**

Castoldi M, Popova AV (2003) Purification of brain tubulin through two cycles of polymerization-depolymerization in a high-molarity buffer. *Protein Expr Purif* 32: 83-88

Inoue Y, Shingyoji C (2007) The roles of noncatalytic ATP binding and ADP binding in the regulation of dynein motile activity in flagella. *Cell Motil Cytoskeleton* 64: 690-704

Kerssemakers JWJ, Munteanu EL, Laan L, Noetzel TL, Janson ME, Dogterom M (2006) Assembly dynamics of microtubules at molecular resolution. *Nature* 442: 709-712

Kinoshita Y, Kambara T, Nishikawa K, Kaya M, Higuchi H (2018) Step sizes and rate constants of single-headed cytoplasmic dynein measured with optical tweezers. *Sci Rep* 8

Miyazono Y, Hayashi M, Karagiannis P, Harada Y, Tadakuma H (2010) Strain through the neck linker ensures processive runs: a DNA-kinesin hybrid nanomachine study. *EMBO J* 29: 93-106

Rice S, Lin AW, Safer D, Hart CL, Naber N, Carragher BO, Cain SM, Pechatnikova E, Wilson-Kubalek EM, Whittaker M *et al* (1999) A structural change in the kinesin motor protein that drives motility. *Nature* 402: 778-784

Yagi T, Uematsu K, Liu Z, Kamiya R (2009) Identification of dyneins that localize exclusively to the proximal portion of Chlamydomonas flagella. *J Cell Sci* 122: 1306-1314

Yasunaga T, Wakabayashi T (1996) Extensible and object-oriented system Eos supplies a new environment for image analysis of electron micrographs of macromolecules. *J Struct Biol* 116: 155-160

### Supplementary Table

**Table S1.** DNA oligonucleotide sequences for Rod. The asterisk indicates the C6dT modified oligonucleotide. Small letter "tttt" indicates the four thymine (4T) linker added to prevent the aggregation of DNA rod.

| Seq Name | Sequence |
|---|---|
| Rod2_001_4T2 | ttttGGGCGATGGCCGTCTATCAtttt |
| Rod2_002 | CCAACGTCGTAATCGCGTGCCTGTTCTTCTCGTCGGAATAAG |
| Rod2_003 | AAAGAACGTGGACTCAAATCAAGTTTTACTATCGATTTTGAGATTAGA |
| Rod2_004 | CACTATTTTCCTGTACCGGGGGTTTCTGCCAGAGCACATCCT |
| Rod2_005 | AAGAGTCGAGGTGCCGTAAAGTGCCTGAAAATGGA |
| Rod2_006_4T2 | ttttTTGAGGATCCCCGGGTGTGAGCCTCCTCACAGtttt |
| Rod2_007 | ACCGAGCTCGAATTCAAAGGGCTTGCAGCAACAAC |
| Rod2_008 | GGCGCTTTCCTTAGTGACCTCCGGCCAGCACATGGTCATAGCTGT |
| Rod2_009_4T | ttttCGACAGTATCGGCCTCAGGAAGAAAACGATTTTTCGTCGCGTCC |
| Rod2_010_4T2 | ttttGCAAACGCGGAAACACCAGAAtttt |
| Rod2_011 | CTGGCAGTGAAGGGTAAAGTTTCGCACTGCCCGAA |
| Rod2_012_L-NdT3_T30 | *T*T*ttttttttttttttttttttttttttttttAGCCGCACAGGCGGCCTGGCACCGCTTCTGGGTCACGTTGGTGTAGATG |
| Rod2_013_4T2 | ttttATATATTCGGTCCTGACGAGTCCGTTGCTGATTGCCGTTCCGtttt |
| Rod2_014 | CATAACGCGGTCATGTGAAATTGTTATCTTGGAAC |
| Rod2_015 | CAACAGGCAACGCAGTCAAATCACCATCGGGATAGTGCCGGAAAAA |
| Rod2_016 | CCAGCCAGCATCGTAACCGTGTTCAAAA |
| Rod2_017_4T2 | ttttAGTAATGTGTAGGTAAAGACATCTGCCAGTTTGAGGGGACGAtttt |
| Rod2_018 | GGAGACAAGGATAAAAATTTTTAGAACCGGGTGAGAAAGGCC |
| Rod2_019_4T2 | ttttGTTTTAATTATGCAATGCCTGtttt |
| Rod2_020 | CTCATATATTTTAACGAGCTTAGACTTCGAGATGG |
| Rod2_021 | CAAAGCGGCAAGGCCTGAACAATAAAAACAGGGAACTAATGCAGAACGC |
| Rod2_022 | AACCAGACCGGAAGAAGATTATAATCATGCTGCTC |
| Rod2_023_4T2 | ttttCGAGTAGTAAATTGGGCTTAAATATCGCtttt |
| Rod2_024 | GTTTGCCGACTCCTCAAGAGATCAACTTAGAGGAA |
| Rod2_025 | TTTAATTAGGATTAAGCGACAGTCCAGACGACGACTCGCTAT |
| Rod2_026 | TGTGAATAGGCTGATTTAGCGGACAAAAAATAGATAAGTCCT |
| Rod2_027 | TACCTTACGGATTGCATCAAACAAACTCCCAGTAGTTGAGCG |
| Rod2_028 | GCTTGCCGCTGAGGCGAAAAACCCACTACGTGAACATCCAGAAACGCTC |
| Rod2_029 | ATTAAATGCCGTCAATAGATAATGGAAAACAATGAGGAGTTAATTCAGT |
| Rod2_030 | ACTAACAACTAATACGCTCAATTGATACGGGATCGATCAACGTAACAAA |
| Rod2_031 | AGAAGCCTTTATTTTCAGGATAGCAAAGTGCGATTTACCCAA |
| Rod2_032 | AAGGCCGCTTTTGCCGATAGTCCGGAATAAACAGT |
| Rod2_033 | TCACCCTCAGCAGCATTTCTTACTCAGGTTGAATT |
| Rod2_034_4T2 | ttttCCATTGCAACACGCATAACCGtttt |
| Rod2_035_4T2 | ttttACCAGGCGGATAAGTGCGGGGTTTTGCTCAGTtttt |

| | |
|---|---|
| Rod2_036 | CCGTCGACATCGCCCAGGAAAACAATATTACCGCCAG |
| Rod2_037 | GAGGGTTATGTGAGTAATTAATTTTCCCTAATTCTGAATCAAACCATTA |
| Rod2_038 | ATACATTATTCGACAACTCGTTCAATATGATATAA |
| Rod2_039 | GTATAGCTGCGCCGTACCTACGCCTTGCTGGTAATCATCACC |
| Rod2_040 | TCACCGTAAACAGCTCGTCTGGTAGAAGAACTCAATGGGGTC |
| Rod2_041_4T | ttttGGAGAATTAACTGAACACCCGGAAACAATCAGTGGATTAG |
| Rod2_042 | CCTTTGCCCGAACGTTAATGGAGGTGTATATTAAG |
| Rod2_043 | AAATAGCCTAATATCAGAGAGAGCCAGCGTAGCGCATGAAAG |
| Rod2_044_4T2 | ttttTCGATAGCAGCACCGTGTCACCAATGAAACCAtttt |
| Rod2_045 | GCCTGTTGAATAACAAGTCAGAGGGTAACACCATT |
| Rod2_046 | CCTTGAAAACATAGAAGTACCTCAGACTAAAATCA |
| Rod2_047_4T2 | ttttCATGTTCAGGCGCATTAGACGtttt |
| Rod2_048 | AGCCTTTACAGAGATATCAACGGTAAAGTTAGAATACATAAA |
| Rod2_049_4T2 | ttttTGTAAATCGAATAAACAAtttt |
| Rod2_050_4T2 | ttttTTAGACTTTCCTTGCTTCtttt |
| Rod2_051_4T | TGAATAAACAAACATGAGGATTTAGAAGTAtttt |
| Rod2_052 | AACGTCAAAAATGAGAACAAGGAATATACGATAGCACCTTTTTTATTAA |
| Rod2_053 | ACCCTGAATTATACTGACAAGAGCATCGAGTGTTGTTCCAGT |
| Rod2_054 | ATATGTAGAAGAATGCGAATTCCACACAACAAGGGTTG |
| Rod2_055 | CCGAGATTACGAGCTGGTGCTGCGGCCACGTCAGCGTAATCT |
| Rod2_056 | GAATAGCCCGATTTAGAGCTTAACCGTTAGTAATA |
| Rod2_057 | CAACTGTCAGTTGGTGGTCTGCACTCTGCGGAAGCATAAAGTATCAAAA |
| Rod2_058 | TTCAGAAAAAACGAGACCAGGACTAAAGAAAATCCCTTATAA |
| Rod2_059 | TGCCAAGAATCAGTGTCCTGGGGTGCCTAATAATCGGC |
| Rod2_060 | CGCTCACGCGGGCCGTTTTCAGAACGTGTATTCATTTAAGAA |
| Rod2_061 | GTAAAGCACTGCGCGCCTGTGGTCAGCAGCTGGCT |
| Rod2_062 | GTTCCGAGAGTGAGCAGACGATCCAGCGGCCAACG |
| Rod2_063 | ATCAAGAGTGGTGCGCGGTTGTGTACATCGAC |
| Rod2_064 | ATGAACGGCAGCACGGATCAAACTTAAATTTC |
| Rod2_065 | AACCGGACCGGACTAAAAAATCCCGTAAAACCAGG |
| Rod2_066 | TACCTTTTTGCGGGTATTCAACCGTTCTAAACGGCCCATTCGCCATTCA |
| Rod2_067 | CGCATAGGCAACCGTCATTTGCCGCCAGTGGGAAG |
| Rod2_068 | TGGCTTACTAAATCGCTATTTTTGAGAGAAATGTGGGTGCGGGCCTCTT |
| Rod2_069 | GACGTTGAAAATCAGGCTGCGTTCTCCGTGGGAAC |
| Rod2_070 | CCCTTATATCACCGATAAGAGCATTATGAAATTAATGCCGGACGTCGGA |
| Rod2_071 | CTGGCTCCTATTATCAAAGCGGGATTGACCGTAATAATATGA |
| Rod2_072 | AGCTGATACCCTGTAATACTTAATTGCT |
| Rod2_073 | CGTTAATAACGAGAGGCGATCAGCGAGTAACAACCGAGGGTA |
| Rod2_074 | GGTTGTACCAAAAAGTCATTTTAAATCAGGAAGAAGACCTTCACGGCTA |
| Rod2_075 | CGCTATTCTTTCATCAACATTATCTACACTCAGAGCATAAAGGAGCTTA |
| Rod2_076 | AGTCAGATAGAGAGGAATTAGATAACCC |
| Rod2_077 | GGTCTTTCCTTTTGTCACCGAAAGCCCAATAAGAAACGATTTATCCTAA |
| Rod2_078 | ATGACCATTGCGGAATTAAAGAAACAAT |

| | |
|---|---|
| Rod2_079 | CAGTCAGCCTATTTCATAGCCCAGAGGCTTTACGAGCATGTAAATCCAA |
| Rod2_080 | AAATCTAATAAACATGCCATCAACGCCAATCAATAATCGGCT |
| Rod2_081 | GTGTACAACTAACGGCCTTGAATCACCGTTGAGAACTTATCATTCCAAG |
| Rod2_082 | ATCTAAAATATCTTTTATTTAGAGGTGAGAAAGAC |
| Rod2_083 | GAACGAGATCAGCTCACCCTCAATTACAAGTCAATAGTGAATATTTAGG |
| Rod2_084 | TGAAAGGAATTGAGCCAGTCATCGGTTTGGTAGCA |
| Rod2_085 | CAGAGGCAGCCTTTCAGAACCAAACATC |
| Rod2_086 | GTCAGTTGGCAAATAAAGGGACAAAAGGTTTGAGG |
| Rod2_087 | ACTTTTTCAAAAAACACCACCAGCAAAACCTTTTTAACCTCCGGCTTAA |
| Rod2_088 | GTACCGCTGCTTTCCATTGGCGTAATAACATCACTCACTAAA |
| Rod2_089 | CCACCCTAATTGTACACGACCGTAGCAATACTTCTGGAGCCC |
| Rod2_090 | TCAGAGCAAGGCTCCATTCTGCCATCACGCAAATTGACGGGG |
| Rod2_091 | CTGAAACGTTTTCAAATAAGAAAAAATAATATCCC |
| Rod2_092 | TAGGAGCTTTTAAAAGTTTGAATTTCATAGGTTTATATTATT |
| Rod2_093 | TTTGTTTACAAGAATTGAGTTCTTGAGCTTTCGGTCGGAACC |
| Rod2_094 | GAAGGTTTATCATTTTGCGGACAAAATTAGAACCGCCCCCTG |
| Rod2_095 | ATAATAAGAGCAAGGTGAATTTAGCGTTGTTAATG |
| Rod2_096 | CAACAGTAACCACCAGAAGGAATGAAACGCCACCCTGCCCGT |
| Rod2_097 | AACAGCCGAAATAGCAATAGCGACGGAAAATCAAAGTAACAG |
| Rod2_098 | TTAGATTAAGACGCAGCCAGTTCGGCATCATTTGG |
| Rod2_099 | AATCATAGGTCTGAATTTAACTTTTCATATTATTC |
| Rod2_100 | AATGCTTTAAACAGATTGCTGAAATATTTATCTTA |
| Rod2_101 | ATATTATTTATCCCGAAACCAACATGTATTATCAAAAGAAAAACAAAGA |
| Rod2_102 | CCAGTTACAAAATAGTCTTTCTCGCCATGAGACTAGAAGATGGCGGAAT |
| Rod2_103 | ATTTTCGTGAGAAGTTTAACAGTAACAT |
| Rod2_104 | TATCATCATATTCCTTACCTGCTCATTTGGTCAGTGAACAAC |
| Rod2_105 | AAAGCCGGCGAACGAGTCTGTGCCAACAATATCTG |
| Rod2_106 | TCGGAACCCTAAAGTTGATTAAGATTCA |
| Rod2_107 | GATGGTGAAAGGAAGGGAAGAAGTGAGGCTGACCT |
| Rod2_108 | ACGCCAGAAACAGCCGTCGGTCCTGCATCTAACTCACATTAACCTGTTT |
| Rod2_109 | CAGCAGGCGAAAATTTGCGTTCGGTGCCGGTGCCCGGTGCCA |
| Rod2_110 | ATGTGCTCTCACGGCAACCAGCAGCCAGGCGCTCACTGCCCGTTTGCCC |
| Rod2_111 | ACCACATGACGGTCGGCACCAGCGGTCCACGCTGG |
| Rod2_112 | TTGGGTACTCCGTGTGTCCAGCATCAGAGTCGGGAAACCTGTGCAGCAA |
| Rod2_113 | GAGAGTTCGTGCCACAAATCGTTAACGGCATCAGCGGGGTCA |
| Rod2_114 | CTTTCCATGCCGGGTTACCTGCTTACGG |
| Rod2_115 | TTGCAGGTGTTCAGGCTGCATTAATGAA |
| Rod2_116 | ACCAACTTCCCACGAAAAAGAGACGCAGCTGGCGA |
| Rod2_117_R-NdT3_T30 | *T*T*ttttttttttttttttttttttttttttTTGTGAGAGATAGACTTTACGCCAGGGTTTTCCTCATTTTTTAACC |
| Rod2_118 | AATCATACTGGAGGGTGAAGGGATAGCTGCAAGGC |
| Rod2_119 | GAACCAGGGAAGGTAATATAAATAAAGCAAGGCTATCAGGTCTAGCCAG |
| Rod2_120 | CCGCCTCACATTCAATGTTTTAAGAATTGAGAGTCTGGAGCAAATTCGC |

| | |
|---|---|
| Rod2_121 | CCCTCAGAGCGCCAAGTACGGATAAATCGAATCGATGAACGGAATAGGA |
| Rod2_122 | AAAGATTATATTCAAAGGGGGGTCTGGCCTTCCTGATTGCCT |
| Rod2_123 | TAGGAATAATACTGGATTAAGACGCCATCAAAAATAACAAGA |
| Rod2_124 | TAATCGTAAAACTAGCATGTCAAATCAGCCAGTCACGACGTT |
| Rod2_125 | AGCAAAATTAAGCATGCTGTACCCCTCAATTATTAAGGACAGAAGTTTC |
| Rod2_126 | ATACAGGCAAGGCAAAATATGGTCATAACATCAGTCGAACTGAATACGT |
| Rod2_127 | AGCATTAACATCCATGTCTGGAGCGTCC |
| Rod2_128 | GAGCCACAGAAAATATTCCATTAGTAGT |
| Rod2_129 | TTGAATCGCTCAACACCGATTAAGAAAA |
| Rod2_130 | CGGAATCCAACTAAAAGACAAGAACAAA |
| Rod2_131 | ATGCAGATGTTTAGACTGGATAAGTTTCTCATATGACCGAGG |
| Rod2_132 | TTGAAAGCAGGTAGATAAGTTACCGGAAACGCTCAATTAAACCAAGTAC |
| Rod2_133 | AGGGAACTGAGATTTGATACAGCCGCCAATTCTTAATCGAGAACAAGCA |
| Rod2_134 | AGGCGCATCAACTAAGCGTCAACCCTCAAGTATCATTTATTTTCATCGT |
| Rod2_135 | GCCACGCTGAGAGCCTATTAGCAGCGGAACGAAAGCGGAACG |
| Rod2_136 | CAAACCCTCAATCAGAGATAGAAATCTCCATGAGG |
| Rod2_137 | CATTAAATTTTTTCTAGCAAGATTATTCGTTGGGTTATATAAAAAGCCA |
| Rod2_138 | ACCTTGCTGAACCTGAAAGCGATAATAACGGGTAA |
| Rod2_139 | AATGCCATAAAGGAATGTACCTACAAAATAAATGCTGATGCATATACAA |
| Rod2_140 | AAATGAAAAATCTAACAGACAGAACAACCTACGAA |
| Rod2_141 | ACCTAAAGTGAGAACGTCACCTGATTGCTCGCAAGACAAAGACCTGTTT |
| Rod2_142 | TCAGGGAACGTTGAAACCCTTCCACCGAGTAAAAGTGGCGAG |
| Rod2_143 | GGAACCCATTGCGATAAGAATGTGTTTTTATAATCAAGCGAA |
| Rod2_144 | TGAGTTTTAGAAAGATATTTTGGTACGCCAGAATCGGCGCT |
| Rod2_145 | ACTACAAACAGTTTTCTTTAAAAAGGGATTTTAGATCACGCT |
| Rod2_146 | TAATTTGCCGAAGCCCTTTTTGAGGGAGAGCCACCTTAACGG |
| Rod2_147 | CAAATATCAGATGATGGCAATGAGGCGACCCAATAACTGGTA |
| Rod2_148 | AACGAGCGTAAGCAGATAGCCAAGGGCGCCTCAGAGGAGTGT |
| Rod2_149 | AAGCATCTATAATCCTGATTGACCAAGTGTAACACCTTTTGA |
| Rod2_150 | ATCCTGAGTTACCAGAAGGAAGTTTACCAACCGCCTACATGG |
| Rod2_151 | CAGCAGCTATACTTCTGAATAATTCGCCAGTACAATCCAGTA |
| Rod2_152 | TTTGCACAAACGCAATAATAAAATCAATCACCCTCTTACCGT |
| Rod2_153 | GTCTTTCCAGAGCCAACGGGTACAGTAGGGCTTAGATTTCAATGATTAT |
| Rod2_154 | ATCTTACCAACGCTCGCACTCCCAGTATCTATATGTCGCGCATCATCAA |
| Rod2_155 | CCAGCTACAATTTTAGCCGTTTATGCGTAATCCAATTTGAATTTTGGAT |
| Rod2_156 | AGGAGCGGGCGCTACTGAGAAACGTGGC |
| Rod2_157 | GGCAAGTGTAGCGGCAGGAACTGAATGG |
| Rod2_158 | CCGCCTGGCCCTGAGCGCGTAACCACCAGCCGATTTGCGCGA |
| Rod2_159 | CCCTTCATCGGCCACCCTTACACTGGTGCGCTTTC |
| Rod2_160 | AGTTTTGACGAGGCATCCGCGCACTCATCTGATTG |
| Rod2_161 | GACGGCCCGCCATGATCCGCCTAAACATACGCGCGGGGAGAGGCAACA G |
| Rod2_162 | GAGACGGGCGGTTTTTCTTTGCTCGTCAGGGCGCG |

| | |
|---|---|
| Rod2_163_4T | CACCAGTGGGCGCGTACTATGCGTGCTTGAACGAACCACCtttt |
| Rod2_164_4T | ttttGGTTTTTCTTTTCCAAGCGGACGTTAGATCTAA |
| Rod2_165 | AACAATCTATGAGCTAAAGGTGCGTATTGGGCGCCAGGGT |
| Rod2_166 | TGCGGCTGGTAATGGGCGGGTCAGTATCATTAACCCT |
| Rod2_167_4T2 | ttttCAACGGAGATTTCTGTTGCCCtttt |
| Rod2_168 | ACCTGCTGCACTCATTTACCAGTCCCGGAAT |
| Rod2_169_4T | ttttGGATAACCTCACCGGAGAGCCGCCAAAATATAGATAC |
| Rod2_170 | CCCCAGCTAAATTGGTTGCGGGGCGAAACGTACAGAGTGCCA |
| Rod2_171 | CATTTGGAAAAGCCCCAAAAATAAACGTAGAGGTG |
| Rod2_172_4T2 | ttttGACGATAAAAACCACGGGAACtttt |
| Rod2_173 | CCACCAGGTTTATTCCAATTCAGGTGGCTGTACCCCGGTTGAAATTCGC |
| Rod2_174 | TAATCAGGGCGCGAGCTGAAATGCGAAC |
| Rod2_175 | GCTTTTGAGCTTTCTAATATTTTGTTAA |
| Rod2_176_4T | ttttGCAAATATTTAAATTGCAGGAAG |
| Rod2_177 | AGGAATTCCAGAGGGTAAAACATTAAATTTTTGTTAATCATA |
| Rod2_178_4T2 | ttttGTCAATAACCTGATTGTATAAtttt |
| Rod2_179 | ATCAATTCTACTAAATAACAGAGTAAAATACATAAACTTAGCAGGCAAA |
| Rod2_180 | TTTAGCTATATTTTTGACCATGCGAGAGACTATCACGCCTGAGATTATA |
| Rod2_181 | TAGTTAATTTCATCTTAAATACCGCCGCCGCAAAGTTTAGTT |
| Rod2_182 | ATACCGACCGTGGGTTGAGGGTGGCAATTTCGCAAATG |
| Rod2_183 | GGGTAATTTGATTCTTGTCACCGGAATAAGATTAG |
| Rod2_184 | CAAAAGAGAGTAGAACACCACCATGATT |
| Rod2_185 | TGCACGTATACAGTATAGTTACATTAAAGAGCAAC |
| Rod2_186_4T2 | ttttATTGGCCTTGATATTCACACGTTTACCAGACtttt |
| Rod2_187 | CCATGTTCGCCAAATCTGAATAGAGCCGATTACTAATTACCGCGCCCAA |
| Rod2_188 | TGTCGAAATAGTAAGCCAGAAACCAGAGAGAATAACAAATCAGATATAG |
| Rod2_189 | AGAATACACTTTCACGCCTGTAGAAACAAAAACTTTTTCAAAAATCATA |
| Rod2_190 | ATTAACACCGCCTGACTGATACTAAACAACTAAAA |
| Rod2_191 | CTTTGACTGTATGGAGCCCTCAACAGTA |
| Rod2_192_4T2 | ttttGTCTTTCCACGAAACAAAGTAtttt |
| Rod2_193 | TTCTGACGATGAATAAAACAGAACAGAGGTGAGGCCCATTAAAATTTTC |
| Rod2_194 | CACAGACGATTTTGGCCCTAAGAGCTAAACAGGAGCACCCGC |
| Rod2_195 | GCGTAACGTAAATGAAATACCTCCTCGTTAGAATCCGCTACA |
| Rod2_196_4T2 | ttttATTTTCAGGTTTAGTTTTGTCtttt |
| Rod2_197 | CAACAGTGGTTAGAACCTACCCATCGGGAGCATTCCGCAGTC |
| Rod2_198 | TAAATCACCCAAAAGAACTGGGGAATAAAACCACCTGGAAAG |
| Rod2_199 | TTGCGGGAAGACTCCTTATTAAAAGAAACAGCATTAAATCCT |
| Rod2_200_4T | AACAAATGACAGGATGATAAAATCCGGTATTCTAAGAACGtttt |
| Rod2_201_4T2 | ttttTACATACATAAAGCAGGTCAGACGtttt |
| Rod2_202 | ACATATACGCAGTAGCGAACCTCCCGACAAGGCTTTAAGGCG |
| Rod2_203_4T2 | ttttCGAGGCGTTTTATGTTAGCAAACGTAGAAAAtttt |
| Rod2_204 | TTGCTATAGGAATCGAAAAAGACGCGAGATAACGGATGGAAG |
| Rod2_205 | AGGTTTTGAAGCCTTAGCAAGACACCGGTATATTTCCTTTTAATATCAA |

| | |
|---|---|
| Rod2_206_4T | AACGTCACTAAATTTAATGGTTTGAAtttt |
| Rod2_207 | CGCGCTTAATGCGCAGAGCGGAACATCGGGTCAGTAATTATT |
| Rod2_208_4T2 | ttttAGCAGAAGATAAAAATAAAGAAATTGCGTAGtttt |
| Rod2_209_4T2 | ttttACGAGCACGTATAAGTTGCTTTGtttt |
